# Pleiotrophin drives a pro-metastatic immune niche within the breast tumor microenvironment

**DOI:** 10.1101/2022.05.02.490334

**Authors:** Debolina Ganguly, Marcel O. Schmidt, Morgan Coleman, Tuong-Vi Ngo, Noah Sorrelle, Adrian TA. Dominguez, Jason E. Toombs, Cheryl Lewis, Yisheng Fang, Fatima Valdes Mora, David Gallego-Ortega, Anton Wellstein, Rolf A. Brekken

## Abstract

Metastatic cancer cells adapt to thrive in secondary organs. To investigate metastatic adaptation, we performed transcriptomic analysis of metastatic and non-metastatic murine breast cancer cells. We found that pleiotrophin (PTN), a neurotrophic cytokine, is a metastasis-associated factor that is expressed highly by aggressive breast cancers. Moreover, elevated PTN in plasma correlated significantly with metastasis and reduced survival of breast cancer patients. Mechanistically, we find that PTN activates NF-kB in cancer cells leading to altered cytokine production, subsequent neutrophil recruitment and an immune suppressive microenvironment. Consequently, inhibition of PTN, pharmacologically or genetically, reduces the accumulation of tumor associated neutrophils and reverts local immune suppression resulting in increased T cell activation and attenuated metastasis. Furthermore, inhibition of PTN significantly enhanced the efficacy of immune checkpoint blockade + chemotherapy in reducing metastatic burden in mice. These findings establish PTN as a previously unrecognized driver of a pro-metastatic immune niche and thus represents a promising therapeutic target for the treatment of metastatic breast cancer.

## Introduction

Metastasis is a highly inefficient process with several rate limiting steps (Cameron et al., 2000; Luzzi et al., 1998). As a result, only a small percentage of cancer cells colonize secondary sites and form successful overt metastases. These ‘metastasis initiating cells’ had to evolve and adapt to overcome the selective pressure of the metastatic cascade (Chambers et al., 2002). Thus, metastatic cancers often demonstrate cellular plasticity (Celia-Terrassa and Kang, 2016) that facilitates evasion of immune surveillance and therapy resistance. Because of enhanced cellular plasticity, metastatic cancer cells can often express developmental proteins that provide survival advantages (Ben-Porath et al., 2008; Lawson et al., 2015). Identifying such key players in the malignant process, especially those that help generate a favorable niche at secondary sites might reveal novel therapeutic targets. Additionally, developmental proteins have restricted expression and function in healthy adults (Warrington et al., 2000), therefore, these factors are often viable therapeutic targets.

Pleiotrophin (PTN), named due to its diverse functions (Li et al., 1990), is an embryonic neurotrophic factor that binds to glycosaminoglycan (GAG) chains (Ryan et al., 2021). It is highly expressed in the central nervous system during the perinatal period and is downregulated thereafter with low expression in adult organs (Wanaka et al., 1993). Physiological functions of PTN include neuronal development (Landgraf et al., 2008; Yanagisawa et al., 2010), adipocyte differentiation (Gu et al., 2007; Sevillano et al., 2019), osteoclast development (Imai et al., 1998), mammary epithelial cell differentiation (Wellstein et al., 1992), angiogenesis (Papadimitriou et al., 2016), response to injury (Himburg et al., 2010; Yeh et al., 1998), and fetal lung development (Weng et al., 2009; Weng and Liu, 2010). Also, PTN is reported to be expressed in inflammatory diseases such as rheumatoid arthritis (Pufe et al., 2003), atherosclerosis (Li et al., 2010) and experimental autoimmune encephalitis (Liu et al., 1998) suggesting it has immune regulatory functions. RNA expression profiling of normal and tumor tissues has shown PTN to be consistently expressed only in glioblastoma (Wang, 2020), however, several studies have indicated PTN is elevated in other cancers such as breast (Ma et al., 2017), lung (Wang and Wang, 2015), and pancreatic cancer (Yao et al., 2017). Even though PTN was discovered in the early 1990’s, the function of PTN remains unexplored in metastatic breast cancer with the majority of earlier studies reporting its pro-angiogenic function in primary tumor development (Papadimitriou et al., 2016).

To study adaptations in metastatic breast cancer, we performed a transcriptomic comparison between highly metastatic and poorly metastatic breast cancer cells. Here, we report PTN to be a ‘metastasis associated factor’ that is enriched at metastatic lesions. We found that PTN expression is associated with advanced disease in breast cancer patients and that elevated expression of PTN might be an independent predictor of metastasis. Additionally, inhibition of PTN significantly reduced lung metastasis in multiple mouse models of breast cancer. Mechanistically, we found PTN activates the NF-kB pathway in cancer cells resulting in elevated expression of cytokines including CXCL5. Inhibition of PTN reverts local immune suppression created by neutrophils resulting in increased activation of CD8 T cells. Furthermore, combining anti-PTN therapy with anti-PD1 and abraxane additively increased the efficacy of immune checkpoint blockade in mice. Overall, our results provide an in vivo functional analysis of PTN in breast cancer progression and demonstrate that inhibition of this previously understudied protein has significant therapeutic potential for the treatment of metastatic breast cancer.

## Results

### Comparison between highly metastatic and poorly metastatic cells reveal distinct metastasis-associated signature

To understand the pathways involved in the metastatic cascade, we performed RNA sequencing on a pair of syngeneic mouse breast cancer cell lines, E0771 and E0771-LG (Figure 1A). E0771 is poorly metastatic and was developed from a spontaneous mouse mammary adenocarcinoma (Ewens et al., 2005; Kitamura et al., 2019; Sugiura and Stock, 1952). E0771-LG cells were derived from a lung metastatic E0771 lesion (Kitamura et al., 2019). E0771 and E0771-LG cells grow similarly as primary tumors (Supplemental Figure 1A), but E0771-LG cells form numerous spontaneous metastatic lung lesions in contrast to parental E0771 cells (Supplemental Figure 1B, C). Bulk RNA sequencing and downstream differential analysis (Figure 1B) allowed us to broadly survey the changes in gene expression involved in various biological processes associated with E0771-LG metastasis. Among the top highly upregulated transcripts in E0771-LG cells, Pleiotrophin (PTN) was a common gene associated with four out of the five top biological gene sets that were different between E0771 and E0771-LG (Figure 1B). PTN, being a neurotrophic cytokine, has been studied in context of neuronal development and has been associated with invasive glioma (Gutmann, 2017; Qin et al., 2017; Shi et al., 2017). Early studies with PTN in breast cancer hinted at potential association of PTN with aggressive disease (Chang et al., 2007). Moreover, Midkine (MDK), a protein related to PTN, was recently reported to be associated with aggressive immune evasive melanoma (Cerezo-Wallis et al., 2020; Olmeda et al., 2017) suggesting that these small heparin binding neurotrophic factors might have previously unexplored important functions in malignancy. To confirm PTN as a metastasis associated factor, we compared PTN expression in the syngeneic cell lines, E0771 and E0771-LG, by RNA fluorescence in situ hybridization (RNA-FISH) (Figure 1C), qPCR (Figure 1D) and western blot analysis (Supplemental Figure 1D). PTN was elevated at the RNA level and at the protein level in metastatic E0771-LG cells. To further test if PTN expression is associated with metastatic cancer cells, we looked at PTN expression in cells derived from MMTV-PyMT mouse genetic model. Cancer cells isolated from metastatic lungs had higher PTN expression compared to cancer cells isolated from primary tumors (Figure 1E). Further, when we compared PTN expression by immunohistochemistry (IHC) in 19 paired tumor and lymph node metastasis samples from human breast cancer patients, we observed higher PTN protein expression in cancer cells in lymph node metastases (Figure 1F, G). These data confirm that elevated PTN expression in cancer cells at the metastatic site is conserved across species irrespective of metastatic site location, i.e., lungs or lymph nodes.

**Figure 1.**
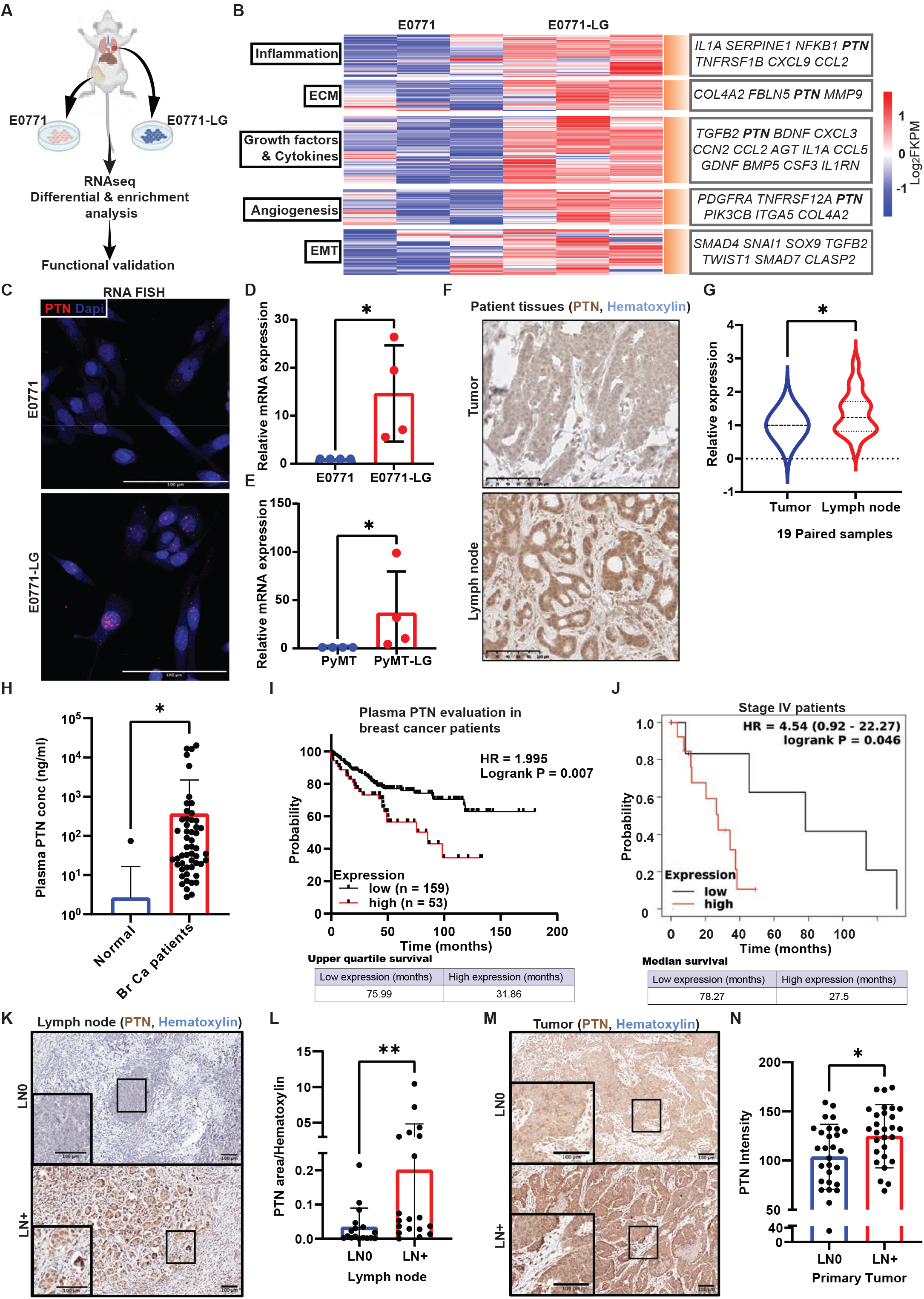
Metastatic cancer cells have enrichment of neurotropic gene Pleiotrophin (PTN) that correlates with poor prognosis in breast cancer patients. **A)** Schematic diagram outlining the strategy for comparison between a highly metastatic lung derived line, E0771-LG, and a poorly metastatic primary tumor derived line, E0771. **B)** Heat map of differentially expressed genes between E0771 and E0771-LG cell lines. Relevant genes that are enriched in E0771-LG cells, in each of the important biological processes, are highlighted in the box on the right. **C)** PTN enrichment in E0771-LG cells was validated by RNA FISH for PTN on cultured cells. **D, E)** PTN enrichment in metastatic lung derived E0771-LG **(D)** and PyMT-LG **(E)** cancer cells were validated by qPCR. Unpaired t-test, *p<0.05. **F)** 19 pairs of breast cancer patient primary tumor and their corresponding lymph node metastasis were stained for PTN by IHC. Representative images are shown at 20x magnification. Scale bar, 100 μm. **G)** Quantification of H is shown as violin plot. Paired t-test, *p<0.05. **H)** PTN expression in plasma^#^ of healthy (no cancer, n = 28) individuals and breast cancer patients (n = 210) were determined by ELISA and is shown as a bar graph. Unpaired t-test, *p<0.05. **I)** Kaplan Meier plot for overall survival of breast cancer patients in (**H**) was generated by comparing survival of patients with high (n = 53) versus low (n = 159) levels of PTN in plasma. Log-rank (Mantel-Cox) test, **p<0.01. **J)** Kaplan Meier plot derived from TCGA database showing overall survival of stage IV breast cancer patients expressing high (n = 13) versus low (n = 7) levels of PTN in their primary tumor. Data were generated from https://kmplot.com/analysis/. **K)** Lymph node tissue from breast cancer patients were obtained^#^ and stained for PTN by IHC. Representative images of PTN staining in lymph nodes from lymph node positive (n = 19) and negative patients (n = 17) are shown at 10x magnification; Images at 20x are shown in insets. Scale bar, 100 μm. **L)** Images from E were quantified in terms of PTN positive area/hematoxylin area. Unpaired t-test, **p<0.01. **M)** Primary tumor tissues^#^ from lymph node positive (n = 29) and negative (n = 29) patients were stained by IHC for PTN. Representative images are shown at 10x magnification and images at 20x are shown in insets. Scale bar, 100 μm. **N)** Quantitation of G is shown as a bar graph of PTN intensity in primary tumors of lymph node negative and positive patients. Unpaired t-test, *p<0.05. ^#^Tissues and plasma samples were obtained from UTSW Tissue Management Shared Resources.

### High PTN expression is associated with aggressive disease and poor survival in breast cancer patients

PTN is a secreted factor and has been reported to be expressed during embryonic development with limited expression in healthy adults (Sorrelle et al., 2017; Wang, 2020). Therefore, we evaluated PTN expression in breast cancer patient plasma. We tested plasma from 28 healthy individuals (no malignancies) and 210 breast cancer patients by ELISA. PTN was detected at a significantly higher level in breast cancer patients (Figure 1H) suggesting it might have potential use as a diagnostic biomarker. Breast cancer patients with plasma PTN concentrations above the detectable level (1 ng/ml) were considered ‘PTN high’ whereas others were considered ‘PTN low’. Interestingly, even though PTN concentration did not significantly differ between Stage IV, lymph node positive and lymph node negative patients (Supplemental Figure 1E), ‘PTN high’ cohort of patients had a worse overall survival than ‘PTN low’ patients (Figure 1I). This suggests that changes in PTN level might be a predictor of prognosis in breast cancer patients. Indeed, patients that had high PTN, regardless of disease stage, had a worse outcome clinically. Concurrently, when we plotted overall survival data of stage IV and lymph node positive breast cancer patients from TCGA, elevated expression of PTN correlated with poor overall survival (Figure 1J, Supplemental Figure 1F). PTN IHC staining of lymph nodes and tumor tissue from patients with lymph node positive (LN+) and lymph node negative (LN-) disease revealed that PTN expression was significantly higher in lymph nodes (Figure 1K, L) as well as in tumors (Figure 1M, N) from LN+ patients. These slides were confirmed, in a blinded fashion, by a breast pathologist (YF). Further, high PTN was associated with lymph node positivity but not with tumor size, age, or hormone status (Supplemental Figure 1G). This suggests that PTN might be an independent predictor of metastasis and poor outcome.

### Cancer cells and stromal components contribute to overall PTN produced within the TME

PTN was first purified from MDA-MB-231 cells, a human breast cancer cell line (Wellstein et al., 1992). We also found that PTN is expressed by cancer cells in human breast tumors. However, the level of PTN in stromal cells in the tumor microenvironment (TME) is unclear. To address this, we accessed single cell RNA sequencing data (scRNAseq) from human breast cancer (Wu et al., 2021) and found that PTN was expressed by a population of tumor cells and lowly by multiple stromal components (Figure 2A, B Supplemental Figure 2A), including endothelial cells. scRNAseq data from late-stage mouse MMTV PyMT tumors also shows PTN is produced by endothelial cells and a sub-population of cancer cells (Supplemental Figure 2B, C, D). We validated the scRNAseq data in patient and mouse MMTV-PyMT tumors by multiplex staining of PTN (by IHC or RNA-FISH) with cancer cell (panCK or PyMT) and endothelial cell (CD31) markers (Figure 2C, D). In MMTV-PyMT lungs bearing metastasis, PTN was enriched at the metastatic lesion whereas little or no PTN was detectable in normal lung (Supplemental Figure 2E). By combining dual FISH and multiplex IHC (Figure 2E a-h), we stained metastatic lesions in MMTV-PyMT lungs for PTN (RNA-FISH), PyMT antigen (cancer cell marker), CD31 (endothelial marker), CD45 (pan-immune cell marker) and podoplanin (fibroblast, lymphatic maker). Multiplex staining revealed that multiple stromal components can contribute to overall PTN that we see in metastatic lesions, however, the majority of PTN is produced by cancer cells. Furthermore, the most studied receptor for PTN, RPTPβ/ζ (Perez-Pinera et al., 2007; Ryan et al., 2016) (Figure 2E i-k), was also enriched in the metastatic lung lesions, suggesting PTN signaling at the metastatic site is potentially functionally relevant.

**Figure 2.**
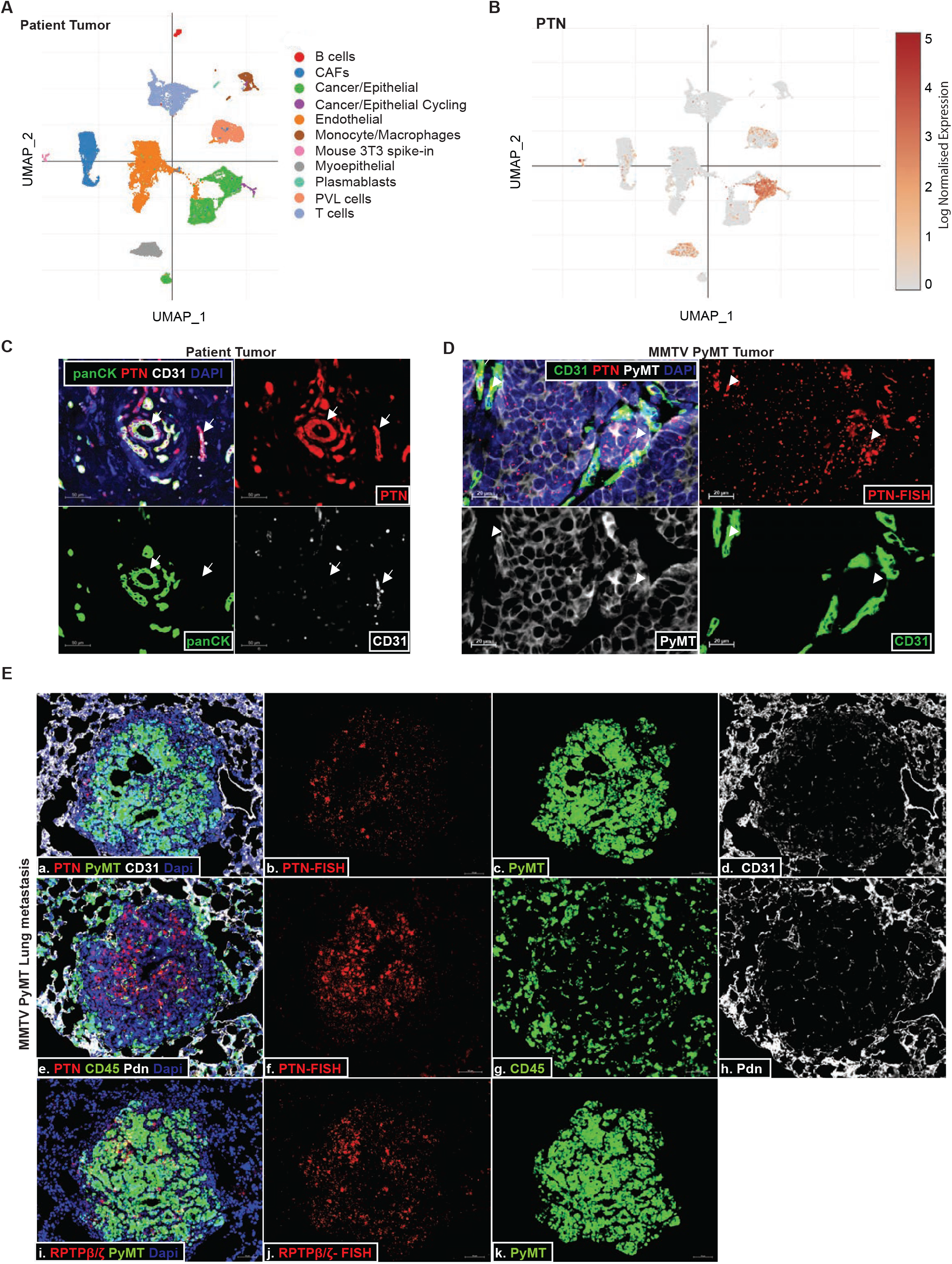
Cancer cells and endothelial cells produce PTN in the TME. **A)** t-distributed stochastic neighbor embedding (tSNE) plot from scRNAseq of a patient’s primary tumor, BC-P1 CID4471, showing the different cell types within the TME. **B)** tSNE plot showing the expression of PTN in different cell types from (A). Data was accessed from the online portal (Wu et al., 2021): https://sinqlecell.broadinstitute.org/. **C)** Breast cancer patient primary tumor FFPE sections were stained by multiplex IHC for PTN (red), panCK (cancer cells; green), CD31 (endothelial cells; white) and DAPI (nucleus; blue). Representative images are shown at 20x magnification. Scale bar, 50 μm. White arrowheads point at co-localization of PTN signal with panCK and CD31. **D)** Representative images of dual RNA FISH of PTN (red) and IHC of CD31 (endothelial cells; green), PyMT (cancer cells; white) in MMTV-PyMT primary tumor are shown at 40x magnification. Scale bar, 20 μm. White arrowheads point at co-localization of PTN signal with PyMT and CD31. **E)** Representative images of dual RNA FISH of PTN (red; a, b), PTN receptor RPTPβ/ζ (red; i, j) and IHC of CD31 (white; a, d), PyMT (green; a, c, I, k), CD45 (green; e, g), podoplanin (white; e, h) in MMTV PyMT lung metastases. Images are shown at 20x magnification. Scale bar, 50 μm.

### Pleiotrophin depletion reduces metastasis in multiple preclinical models of breast cancer

Since PTN can be produced by cancer cells and stromal components, we generated *Ptn-null* MMTV-PyMT FVB mice to study the function of PTN in breast tumor progression. We compared *WT* MMTV PyMT and *Ptn-null* MMTV-PyMT tumor histology of at 8, 10, 13, and 16 weeks of age. *WT* MMTV PyMT tumors formed intraepithelial neoplasia by 8 weeks, early carcinoma by 10 weeks and late carcinoma with widespread necrosis by 13 weeks (Figure 3A), similar to previously described studies (Attalla et al., 2021; Lin et al., 2003). *Ptn-null* MMTV-PyMT tumors, however, displayed slower progression (Figure 3A) showing hyperplasia by 8 weeks, intraepithelial neoplasia by 10 weeks, early carcinoma by 13 weeks and late carcinoma by 16 weeks. This phenotype is complementary to aggressive scirrhous tumors observed in PTN over-expressing MMTV-PyMT-*Ptn mice* reported previously (Chang et al., 2007). Slower tumor progression translated into increased overall survival of *Ptn-null* mice compared to *WT* MMTV-PyMT mice (Supplemental Figure 3A). Alongside, *Ptn-null* mice had smaller tumors at 10 and 13 weeks (Figure 3B, C) as well as fewer metastasis to lungs (Figure 3D, E) at 13 weeks.

**Figure 3.**
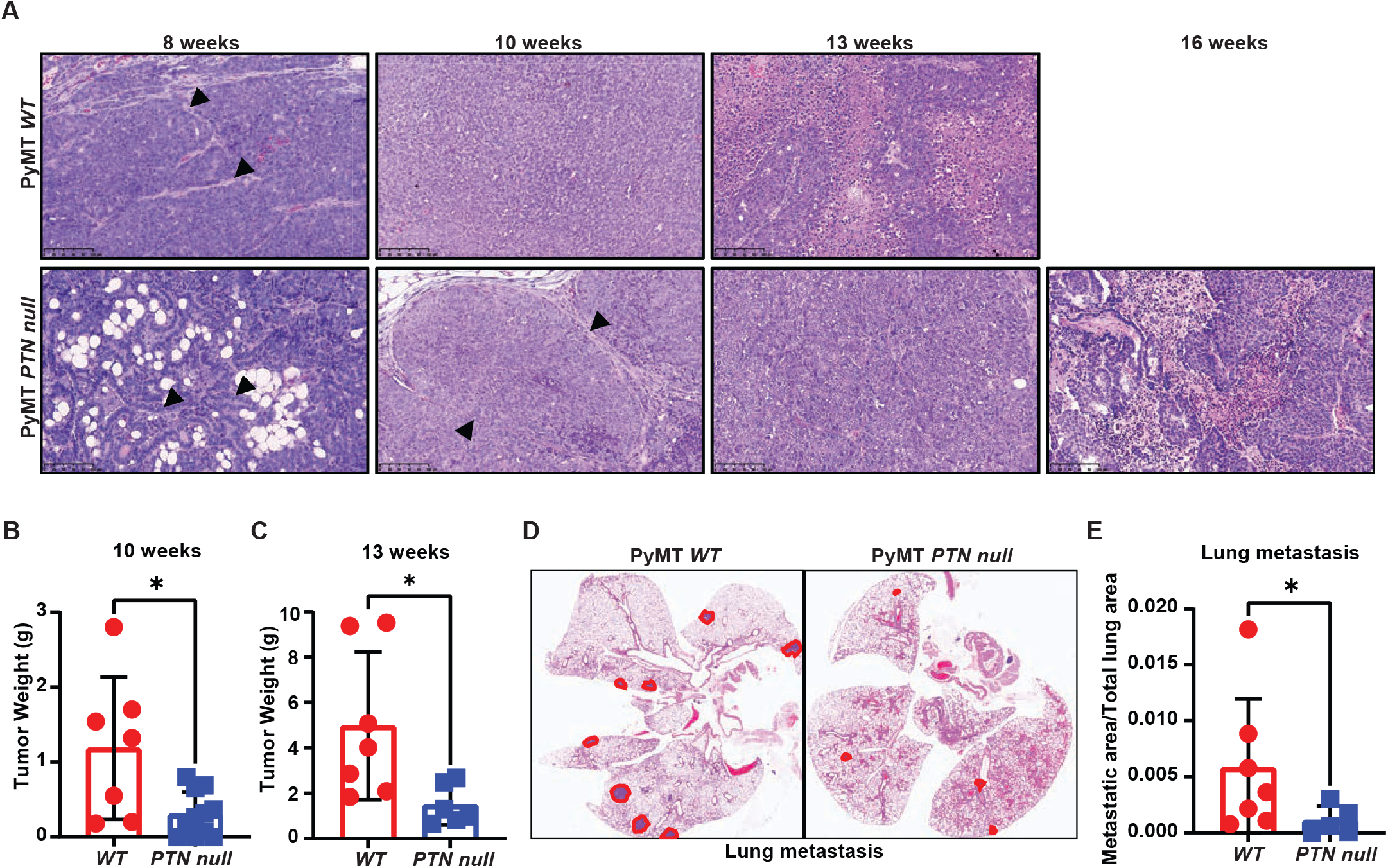
Genetic ablation of PTN reduces breast cancer progression and metastasis in mice. **A)** Tumor progression in *WT* and *PTN null* mice was evaluated by examining tumor histology at 8, 10, 13 and 16 weeks. Representative H&E images of MMTV-PyMT primary tumor from *WT* or *PTN null* mice are shown at 20x magnification. Scale bar, 100 μm. Arrows in 8 weeks point at myoepithelial layer surrounding intraepithelial neoplasia in WT and at normal mammary epithelial cells in PTN null mammary sections. Arrows in 10 weeks point at myoepithelial layer surrounding intraepithelial neoplasia in PTN null mammary tumor sections. **B, C)** Tumor weight in MMTV PyMT *WT* and *PTN null* mice at 10 weeks **(B)** & 13 weeks **(C)** of age. Unpaired t-test, *p<0.05. **D)** Representative H&E images of lung metastasis in MMTV-PyMT *WT* and *PTN* null mice at the end of 13 weeks. Area of metastasis are circled in red. **E)** Lung metastatic burden from (D) was quantitated in terms of metastatic area/total lung area. Unpaired t-test, *p<0.05.

To delineate functional relevance of PTN in tumor initiation versus tumor progression, we utilized a mouse monoclonal antibody that has been reported to neutralize PTN activity, 3B10 (patent: US 20140294845A1). We tested the specificity of 3B10 for PTN by ELISA (Supplemental Figure 3B) and confirmed that it only binds to PTN and not to structurally similar protein MDK or other small heparin binding proteins such as fibroblast growth factor (FGF). We used 3B10 to determine whether PTN is important for progression of established breast tumors by exploiting two orthotopic triple negative breast cancer models, 4T1 (Figure 4A-E) and E0771-LG (Figure 4I, J). Tumor-bearing mice with established and palpable (~50 mm^3^) tumors were treated with an IgG control antibody, C44, or 3B10. In each of these aggressive models, 3B10 treatment did not change primary tumor growth (Figure 4A, I), however, 3B10 did reduce metastasis to lungs (Figure 4B-C, J). In the 4T1 model, we also observed more than 50% reduction in the incidence of lymph node metastasis (Figure 4D, E). This phenotype held true in the MMTV PyMT spontaneous breast cancer model as well. In *WT* MMTV-PyMT mice therapy was initiated after 8 weeks of age, once tumors had already formed, again 3B10 treatment did not reduce primary tumor burden (Figure 4F) but inhibited metastasis to lungs (Figure 4G, H). We also used previously characterized cells derived from MMTV PyMT tumors, Met-1, in an orthotopic setting. Here 3B10 slowed primary Met-1 tumor growth. Met-1 cells are slow growers in vivo and poorly metastatic; we did not see evidence of lung or lymph node metastases in control or 3B10 treated animals bearing Met-1 tumors (Supplemental Figure 3C, D).

**Figure 4.**
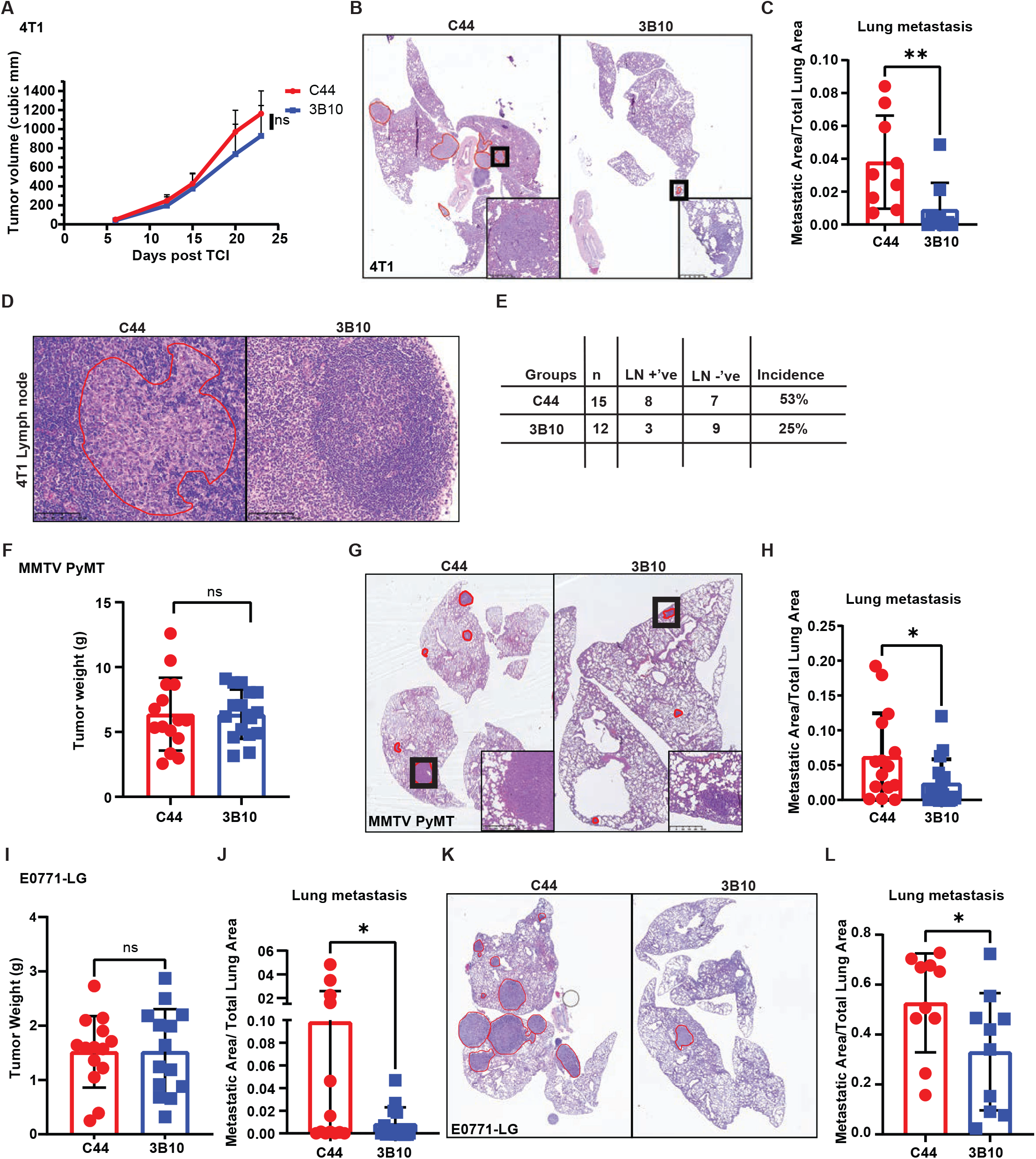
Pharmacological inhibition of PTN (3B10, mAb) reduces metastasis in multiple preclinical mouse models of breast cancer. **A)** Tumor growth curve of orthotopically implanted 4T1 tumors in *WT* BALB/c mice treated with IgG control (C44) or anti-PTN therapy (3B10). Treatment started once tumors were palpable (~50 mm^3^). **B)** Representative H&E images of lung metastasis from 4T1 tumor bearing mice treated with C44 or 3B10. Area of metastasis are circled in red. Insets show metastatic lesions in lungs at 10x magnification. Scale bar, 250 μm. **C)** Metastatic area/total lung area from (B). Unpaired t-test, *p<0.05. **D)** Representation of H&E images of lymph node metastasis in mice from A. Area of metastasis are circled in red. Images shown at 20x magnification. Scale bar, 100 μm. **E)** Quantitation of incidence of metastasis to lymph nodes from (D) by H&E evaluation. **F)** Tumor weight in MMTV-PyMT mice treated with C44 or 3B10 at the end of 12 weeks of age. Treatment started when mice were 8 weeks of age. **G)** Representative H&E images of lung metastasis from MMTV-PyMT mice treated with C44 or 3B10. Area of metastasis are circled in red. Insets show metastatic lesions in lungs at 10x magnification. Scale bar, 250 μm. **H)** Metastatic area/total lung area from (G). Unpaired t-test, *p<0.05. **I)** Tumor weight of mice bearing E0771-LG tumors treated with C44 or 3B10. Treatment started once tumors were palpable (~50 mm^3^). **J)** Metastatic area/total lung area of (I). Unpaired t-test, *p<0.05. **K)** Representative H&E images of metastatic lungs from experimental metastasis study using E0771LG cells in WT C57BL/6 mice that were treated with C44 or 3B10. Treatment started a day prior to tumor cell injection. Area of metastasis are circled in red. **L)** Metastatic area/total lung area from (K). Unpaired t-test, *p<0.05.

To determine if PTN is relevant in cancer cell seeding and colonization of secondary sites, we injected E0771-LG cells via the tail vein. 3B10 or C44 therapy was initiated a day prior to tumor cell injection and mice were thereafter dosed twice weekly. 3B10 treatment significantly reduced metastasis (Figure 4K, L) suggesting PTN is relevant in colonization of secondary sites.

### PTN facilitates inflammation, particularly neutrophil recruitment, by activating the NF-kB pathway in cancer cells

To uncover pathways that PTN influences to promote metastasis, we took an unbiased approach (Figure 5A) through bulk RNA sequencing. 4T1 and MMTV-PyMT tumors harvested from mice treated with C44 or 3B10 were subjected to RNA sequencing. Additionally, we generated PTN knockdown E0771 (E0771shPTN, Supplemental Figure 4A, B) and E0771-LG cells (E0771-LGshPTN, Supplemental Figure 4C, D) and performed RNA sequencing from three replicates each. Amongst the differentially expressed genes (DEGs), NF-kB mediated inflammatory genes (Figure 5B), especially cytokines, were downregulated with PTN knockdown. We then combined GSEA from all 4 experiments and looked for biological processes that were altered by PTN depletion or blocking. Only interferon and inflammation response related transcripts (Figure 5C) were universally reduced in all scenarios. Further, we then took the top 3000 upregulated genes (p<0.05) in control tumors or cells from each of the 4 experiments and ran a motif analysis in Metascape. Using their TRRUST algorithm (Han et al., 2018), we determined potential transcription factors that could control DEGs in each of the 4 experiments. Only 2 transcription factors were common amongst all, namely NF-kB1 and Jun (Figure 5D). To validate PTN mediated NF-kB activation, we stimulated E0771 cells with recombinant mouse PTN in presence or absence of 3B10.

**Figure 5.**
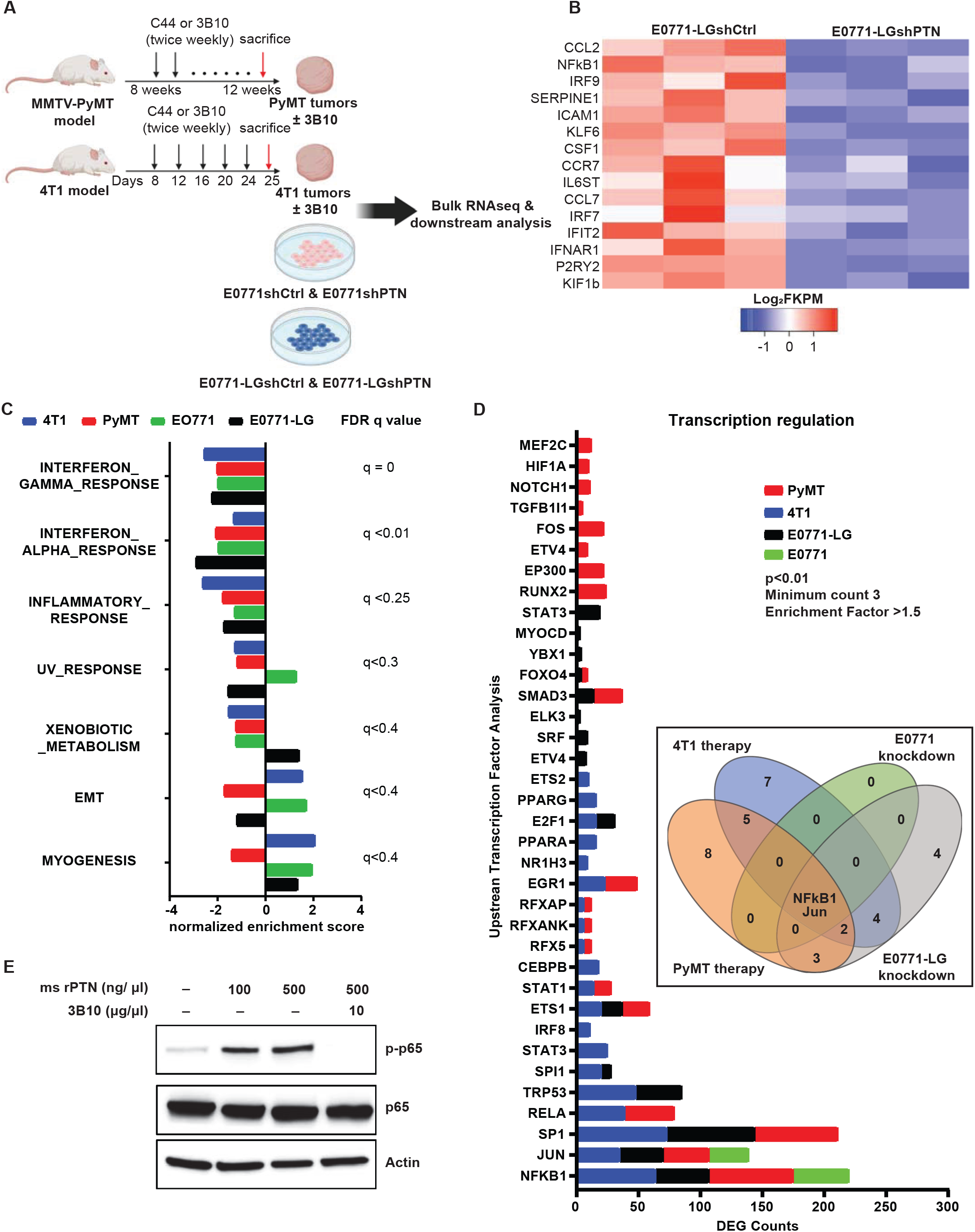
PTN promotes inflammation by augmenting NF-kB activity in cancer cells. **A)** Schematic outlining the RNA sequencing strategy to investigate the effect of PTN depletion in 4 different experiments. **B)** Heat map of differentially expressed inflammation related genes between E0771-LGshCtrl and E0771-LGshPTN cells. **C)** Bar graph of normalized enrichment score (NES) obtained from gene set enrichment analysis (GSEA) of RNA sequencing data as shown in A. Highest q value amongst the 4 experiments is shown on the right. **D)** Bar graph of potential upstream transcription factors regulating DEGs in each of the 4 experiments in A. Inset displays a Venn diagram showing NFκB and Jun are the only two common potential transcription factors that can regulate the DEGs from A as determined by TRRUST analysis using metascape. **E)** Western blot analysis of phospho p65 and total p65 in E0771 cells stimulated with recombinant mouse PTN (100 ng/μl or 500 ng/μl) in presence or absence of 3B10 for 30 minutes.

PTN stimulated phosphorylation of p65, an effect abolished by 3B10 (Figure 5E). Further, many chemokines that are NF-kB target genes and are important in immune cell trafficking were altered by PTN depletion. Most notably, neutrophil associated genes were downregulated with 3B10 treatment (Figure 6A). A potent neutrophil recruiting cytokine and NF-kB downstream target gene (Chao et al., 2016; Deng et al., 2021), CXCL5, was downregulated at the RNA and protein level with 3B10 treatment in MMTV-PyMT-derived tumors (Figure 6A and 6B, Supplemental Figure 5A). This was accompanied by a reduction in neutrophils (Ly6G+ cells) within the TME as determined by IHC (Figure 6C, D). These results were validated in the 4T1 model. CIBERSORT analysis of RNAseq data in the 4T1 model revealed a reduction of neutrophils in 4T1 tumors from mice treated with 3B10 (Figure 6E). Complementing the reduction in Ly6G+ cells, we saw a decrease in S100A9 + cells, a neutrophil & MDSC marker (Averill et al., 2011), within MMTV-PyMT and 4T1 tumors (Supplementary Figure 5B, C, D). Similar to primary tumors, we observed that 3B10 treatment resulted in a reduction in overall CXCL5 expression in 4T1 metastatic lung lesions (Figure 6F). This translated to an overall decrease in neutrophils (Ly6G + cells) within lungs, as determined by IHC and flow cytometry, (Figure 6G, H Supplementary Figure 5E) in the 3B10 treatment group. Neutrophils have been associated with tumor metastasis in breast cancer (Albrengues et al., 2018; Park et al., 2016; Szczerba et al., 2019; Wu et al., 2020; Yang et al., 2020). To determine if neutrophils are required for PTN-mediated metastasis, we performed a Ly6G depletion study in presence or absence of 3B10 (Figure 6I, J, Supplementary Figure 5F, G, H). 3B10 robustly reduced lung metastasis as did depletion of Ly6G^+^cells. However, in the absence of Ly6G+ cells (Supplementary Figure 5F, G), 3B10 did not have an additive effect, supporting our hypothesis that PTN might be enhancing lung metastasis through neutrophil recruitment.

**Figure 6.**
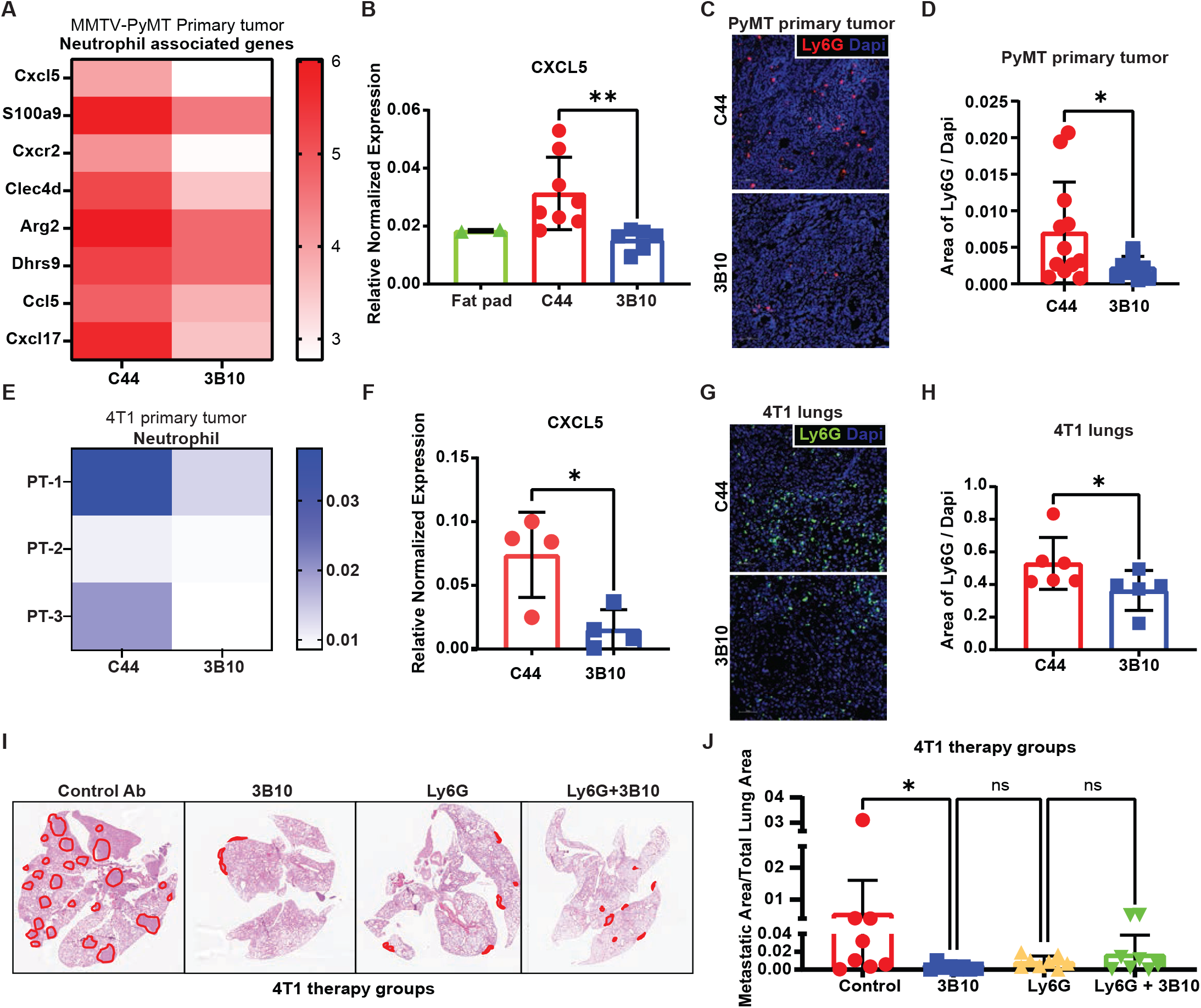
PTN promotes an inflamed tumor microenvironment by recruiting neutrophils. **A)** Heat map highlighting changes in neutrophil associated genes in MMTV PyMT primary tumors treated with C44 or 3B10. **B)** CXCL5 expression in normal mammary fat pad or Met-1 (MMTV-PyMT derived) tumors treated with C44 or 3B10 was determined by cytokine array. Unpaired t-tailed test, **p<0.01 **C)** MMTV-PyMT tumors from mice treated with C44 or 3B10 were stained for neutrophils (Ly6G+ cells) by immunofluorescence (IF). Representative images are shown at 10x magnification. Scale bar, 50 μm. **D)** Quantification of C. Unpaired t-tailed test, *p<0.05. **E)** CIBERSORT analysis using RNA sequencing data of 4T1 tumors show decrease in neutrophil infiltration with 3B10 treatment. Analysis was performed at the online portal: http://timer.cistrome.org/. **F)** CXCL5 expression in 4T1 lungs from mice treated with C44 or 3B10 was determined by cytokine array. Unpaired t-tailed test, *p<0.05. **G)** 4T1 metastatic lungs from mice treated with either C44 or 3B10 were stained for neutrophils (Ly6G+ cells) by IF. Representative images are shown at 10x magnification. Scale bar, 50 μm. **H)** Quantification of Ly6G+ cells from (G). Unpaired t-tailed test, *p<0.05. **I)** Representative H&E images of lung metastases in WT BALB/c mice bearing 4T1 tumors. WT BALB/c mice were treated with either isotype control (IgG), 3B10, Ly6G or Ly6G+3B10. **J)** Quantification of I in terms of metastatic area/total lung area. Kruskal-Wallis test, *p<0.05.

### Blocking PTN reverts immune suppression and has an additive effect in treatment of pre-existing metastases when combined with immune checkpoint blockade and chemotherapy

PTN blockade by 3B10 reduces lung metastasis in multiple models of metastatic breast cancer. Since PTN is enriched in the metastasis microenvironment (Supplemental Figure 2E), we next focused on the changes happening locally in the metastatic lesion. For this, we selected lungs that had metastases from C44 and 3B10 treatment groups and performed multiplex IHC to evaluate the immune landscape. Interestingly, we found that the iNos to Arg1 ratio (Figure 7 A, B) was higher in lungs of 3B10 treated mice. Additionally, 3B10 treatment was associated with a higher number of cytotoxic T cells (Figure 7 A, C) (CD3^+^ CD8^+^ cells) and granzyme B^+^ CD8^+^ T cells (Figure 7 A, D), suggesting an active local anti-tumor immune response. To determine if the anti-metastatic effect of 3B10 was dependent on the adaptive immune system, we implanted 4T1 cells orthotopically in NSG mice and treated with C44 or 3B10. In this setting 3B10 did not reduce 4T1 lung metastasis (Figure 7E, F, Supplementary Figure 5I). These results suggest that PTN production by cancer cells is critical in maintaining a local immune suppressive metastatic niche to prevent T cell mediated tumor cell killing (Figure 8).

**Figure 7.**
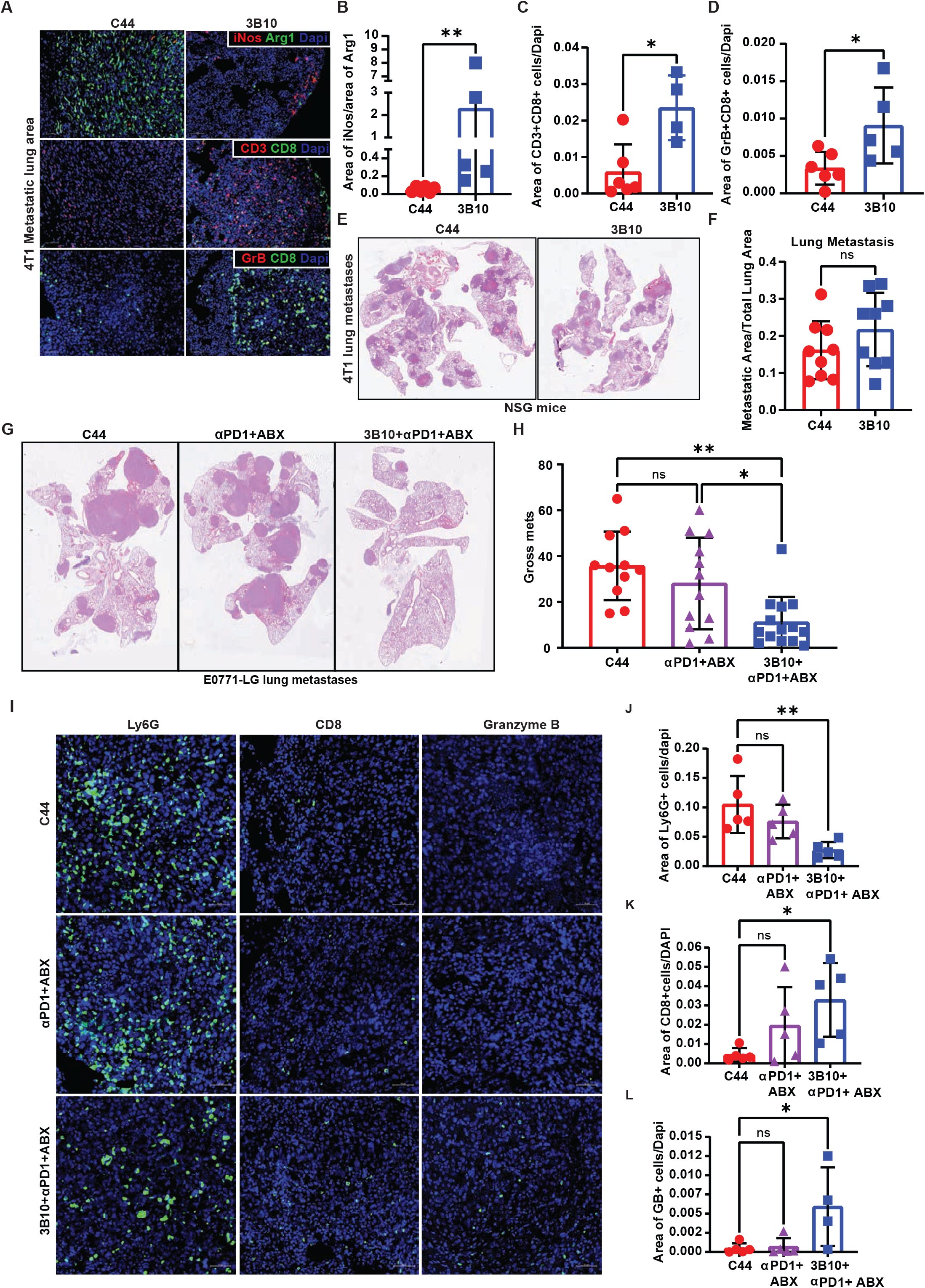
Inhibition of PTN reverts immune suppression and enhances effectiveness of immune checkpoint blockade and chemotherapy. **A)** 4T1 metastatic lungs from mice treated with C44 or 3B10 were stained for pro-inflammatory marker (iNos, red) and immunosuppressive marker (Arg1, green) and cytotoxic T cell markers CD3 (red), CD8 (green), Granzyme B (GrB) by IF. Representative images of IF staining of metastatic lesions in 4T1 lungs are shown at 10x magnification. Scale bar 50 μm. **B-D)** Quantification of A. Unpaired t-test, *p<0.05 **p<0.01. **E)** Representative H&E images of lung metastases in NSG mice bearing 4T1 tumors. NSG mice were treated with either isotype control C44 (IgG) or 3B10. **F)** Quantification of M in terms of metastatic area/total lung area. Unpaired t-test, ns>0.05. **G)** Representative H&E images of metastatic lungs from experimental metastasis study using E0771-LG cells in WT C57BL/6 mice that were treated with C44, anti-PD1+Abraxane or 3B10+anti-PD1+Abraxane. Treatment started on day 7 after tumor cell injection. **H)** Quantification of 0 in terms of gross metastasis. Kruskal-Wallis test, *p<0.05. **I)** Metastatic lungs from G were stained for neutrophils (Ly6G+), CD8 cells and granzyme B+ cells. Representative images of IF staining of metastatic lesions in 4T1 lungs are shown at 10x magnification. Scale bar 50 μm. **J-L)** Quantification of I. Kruskal-Wallis test, **p<0.01, *p<0.05.

**Figure 8.**
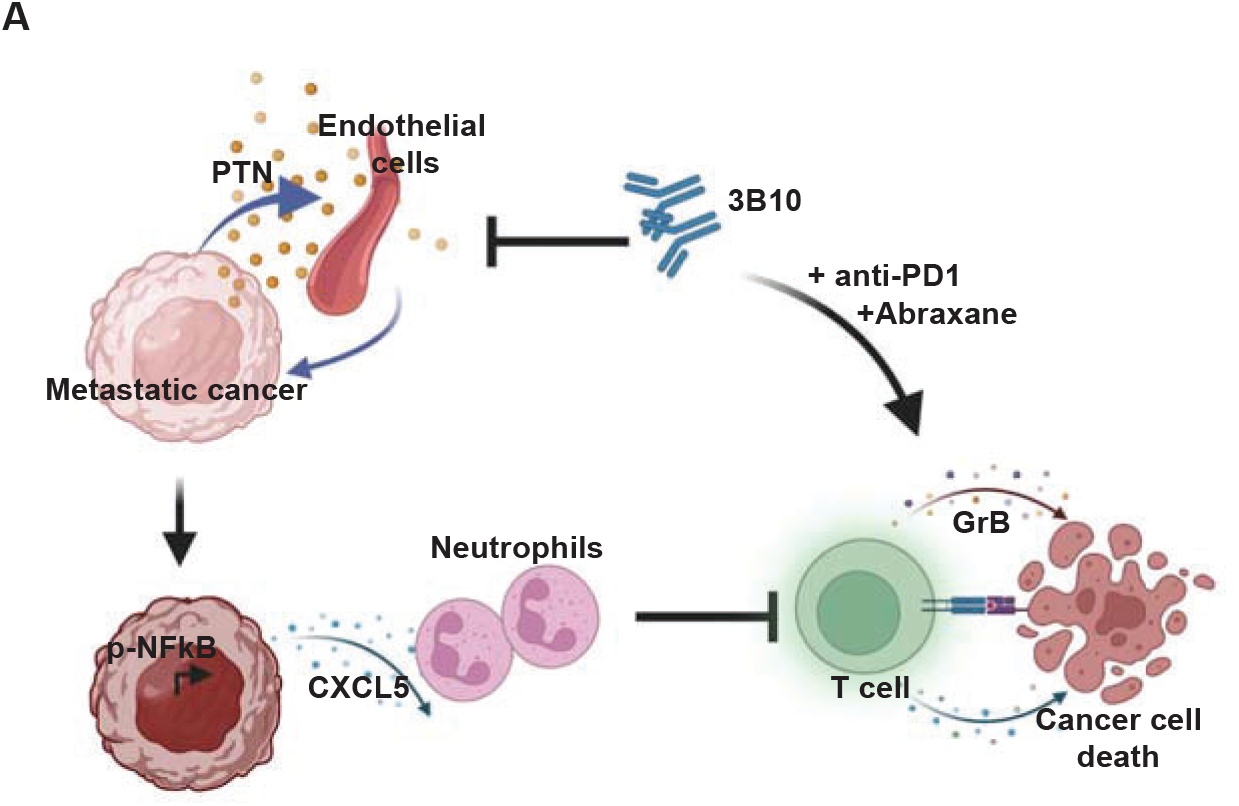
PTN drives a pro-metastatic immune niche. Briefly, the TME of metastatic breast cancer is enriched in cancer cell and endothelial cell derived PTN. PTN promotes an inflamed TME driven by NFκB signaling in cancer cells. PTN-high tumors and lung metastases are rich in immunosuppressive neutrophils that causes T cell dysfunction. Inhibition of PTN reverts immune-suppressive TME thereby increasing the efficacy of checkpoint blockade and chemotherapy in treating pre-existing metastases in mice.

Pembrolizumab in combination with chemotherapy was recently approved by FDA for treatment of metastatic triple negative breast cancer (TNBC) (www.fda.gov/). Given the change in the immune landscape of metastatic lesions after PTN blockade, we speculated that 3B10 combined with immune checkpoint blockade and chemotherapy might show therapeutic efficacy in reducing metastatic lesions. To mimic metastatic TNBC, we injected C57BL/6 female mice with E0771-LG cells via the tail vein and waited for 7 days by which time micro-metastases are formed in lungs (Kitamura et al., 2019). Therapy with IgG control antibody (C44), anti-PD-1 plus abraxane or 3B10 in combination with anti-PD-1 and abraxane was initiated on day 7 post tumor cell injection. While anti-PD-1 plus abraxane showed a mild reduction (not significant) in metastasis compared to the control arm, addition of 3B10 to anti-PD-1 and abraxane significantly reduced lung metastastic burden (Figure 7G, H). Additionally, triple therapy was more effective than 3B10 alone or 3B10 plus anti-PD-1 (Figure 7G, Supplemental Figure 5J). Although the number of PD-1^+^CD8^+^ cells (Supplemental Figure 5L) did not differ amongst the groups, there were significantly lower immune suppressive neutrophils (Ly6G^+^ and Ly6G^+^Arg1^+^ cells) (Figure 7I, J, Supplemental Figure 5K), increased CD8^+^ cells (Figure 7I, K) and granzyme B^+^ cells (Figure 7I, L) in lungs of mice treated with triple combination therapy. These data support that blocking PTN might be clinically promising when combined with immune checkpoint blockade and chemotherapy for the treatment of metastatic breast cancer.

## Discussion

Overall, our results, for the first time, present a comprehensive in vivo functional analysis of PTN (Figure 8) in breast cancer. PTN in the metastatic microenvironment activates the NF-kB pathway in cancer cells leading to increased cytokine production including CXCL5, which results in elevated neutrophil recruitment and contributes to immune suppression. Blocking PTN pharmacologically or genetically reduced metastasis by reverting immune suppression created by neutrophils. This leads to increased activation of CD8 T cells locally at the metastatic niche. Combining anti-PTN with anti-PD1 and abraxane further reduces lung metastasis thus presenting a promising addition to the current regimen for treating metastatic TNBC.

PTN abundance and function in CNS has naturally resulted in it being studied in context of brain tumors. PTN signaling helps maintain glioblastoma stem cells (Shi et al., 2017) and facilitates glioma cell invasion (Qin et al., 2017). Consequently, RNA expression profiling of normal and tumor tissues in the CNS shows PTN to be consistently expressed at higher levels in glioblastoma (Wang, 2020). PTN expression in other cancers, specifically breast cancer has provided mixed results (Garver et al., 1994; Giamanco et al., 2017), with data from TCGA showing lower RNA expression of PTN in breast cancer compared to normal tissue (Wang, 2020). However, a previous clinical study on an Asian cohort of patients showed PTN protein expression to be elevated in breast cancer patients (Ma et al., 2017). This is consistent with our results. Our samples included patients from different stages, subtypes, age, and racial background and represents the largest investigation of PTN in human breast cancer samples to date. A discrepancy between RNA and protein expression levels was also true for a recent study of PTN family member, MDK, in melanoma (Cerezo-Wallis et al., 2020); thus, highlighting the need to evaluate protein levels for these small growth factors. Notably, even though in our study there were not significant differences between Stage IV, LN+ and LN-patient PTN plasma levels, we saw a significant correlation of high PTN expression with metastasis and poor survival. This suggests that irrespective of stage, PTN can predict aggressive disease.

When studied in the context of breast cancer, PTN has been reported to be cancer cell derived factor only (Wellstein et al., 1992). While cancer cells are the major source of PTN in the TME, we found that multiple stromal cells including endothelial cells also contribute to PTN expression in the TME. Though initially studied as a pro-angiogenic factor, 3B10 fails to alter primary tumor growth or microvessel density (data not shown) in TNBC orthotopic mouse models. Instead, we report for the first time, PTN to be a ‘metastasis associated factor’ in mouse and human breast cancer. Interestingly, in mouse lung metastasis models, PTN expression is enriched at the metastatic niche with little or no expression in surrounding normal lungs. This highlights that metastatic colonies create a local niche to facilitate tumor cell survival (Celia-Terrassa and Kang, 2016). It will be important to investigate what controls local PTN expression. A possibility is that PTN expression is induced as part of an essential adaptation process triggered in cancer cells when they colonize a new site. Aligned with this notion, we report that PTN is essential in forming robust metastasis as blocking it reduces metastatic outgrowth in multiple mouse models of breast cancer. It is intriguing that the absence of PTN slows spontaneous primary tumor growth in *Ptn-null* mice but fails to affect primary tumor growth when 3B10 treatment is initiated once tumors are established. This suggests PTN might also contribute to early tumor development. This hypothesis is consistent with a previous report of PTN overexpressing MMTV PyMT-*PTN* tumors showing rapid tumor progression compared to MMTV PyMT *WT* tumors (Chang et al., 2007). Potentially, PTN related adaptations that occur in cancer cells during early tumor development may be conserved during metastatic colonization events. Alternatively, PTN expressing cancer cells may be better equipped to survive the insults of the metastatic cascade. Consistent with these hypotheses, experimental metastasis assays in presence of 3B10 reduces metastasis.

Though expressed during injury and in several auto-immune diseases (Sorrelle et al., 2017), PTN function in immune regulation remains unexplored. Through our unbiased transcriptomic approach, we show that blocking or depletion of PTN consistently resulted in decreased inflammation. We propose that this is mediated in part by inhibition of PTN-mediated activation of NF-kB pathway in cancer cells. Previously it was reported that PTN driven NF-kB activation helps MDA-MB-231 cells escape doxorubicin-induced cell cycle arrest through expression of CDKN1A (Huang et al., 2018). This was mediated through PTN binding to RPTPβ/ζ and inhibiting its phosphatase activity. Consistently, we found enrichment of RPTPβ/ζ in the metastatic niche suggesting NF-kB activation could be occurring through this pathway. Given that PTN is promiscuous in its binding to cell surface receptors (Sorrelle et al., 2017), delineating PTN function through each of its receptors presents a challenge in the field. However, our results implicate NF-kB to be a downstream effector of PTN driven immune regulation. NF-kB activation causes cytokine alteration especially CXCL5 in cancer cells, which leads to neutrophil recruitment. It is noteworthy that by only inhibiting PTN, the metastatic niche shifted to a pro-inflammatory local microenvironment with high iNos to Arg1 ratio, increased CD8 T cell infiltration and increased presence of active effector granzyme B^+^ CD8 cells. Consequently, blocking PTN in combination with anti-PD-1 therapy and abraxane, in mice with established TNBC metastatic disease, had a significant therapeutic benefit especially compared to anti-PD-1 + abraxane alone, which is currently being used to treat metastatic TNBC patients. Similar to our findings, PTN family member MDK (Cerezo-Wallis et al., 2020) was associated with immune checkpoint blockade unresponsive melanoma. In that study, MDK promoted immune suppression through NF-kB driven Arg1 expression in macrophages that mediated T cell dysfunction. These data along with ours present the intriguing concept of blocking PTN and MDK to enhance the efficacy of immune checkpoint blockade.

In summary, our study identifies PTN as metastasis associated factor that can facilitate local immune suppression favoring cancer cell colonization at secondary sites and therefore presents a novel therapeutic target in treatment of metastatic TNBC.

## Acknowledgement

This work was supported by grants from the Mary Kay Foundation, Metavivor, NIH CA243577 and the Effie Marie Cain Fellowship in Angiogenesis Research to RAB. We thank Dr. John Chute for provision of *Ptn-null* mice. We acknowledge helpful input from other members of the Brekken laboratory, especially Drs. Huocong Huang, Yuqing Zhang, Jill Westcott and Raghav Chandra. We also thank Drs. Srinivas Malladi and Maralice Conacci-Sorrell for critical evaluation of the manuscript. Lastly, we thank the following UT Southwestern core facilities: McDermott Center Next-Generation Sequencing Core, Moody Foundation Flow Cytometry Core, Whole brain microscopy facility, Live Cell Imaging core, Histopathology core and Tissue Management Shared Resources (P30 CA142543).

## Author Contributions

DG: study concept and design, acquisition of data, analysis, interpretation of data and drafting of the manuscript. MS: acquisition of data and bioinformatic analyses. MC: acquisition of data. TN: acquisition of data. NS: acquisition of data. AD: acquisition of data. JT: acquisition of data. CL: Human sample selection. YF: pathological scoring of human samples. FM: acquisition of data. DO: acquisition of data. AW: interpretation of data. RAB: study concept and design, interpretation of data, and drafting of the manuscript.

---

**Supplemental Figure 1.**
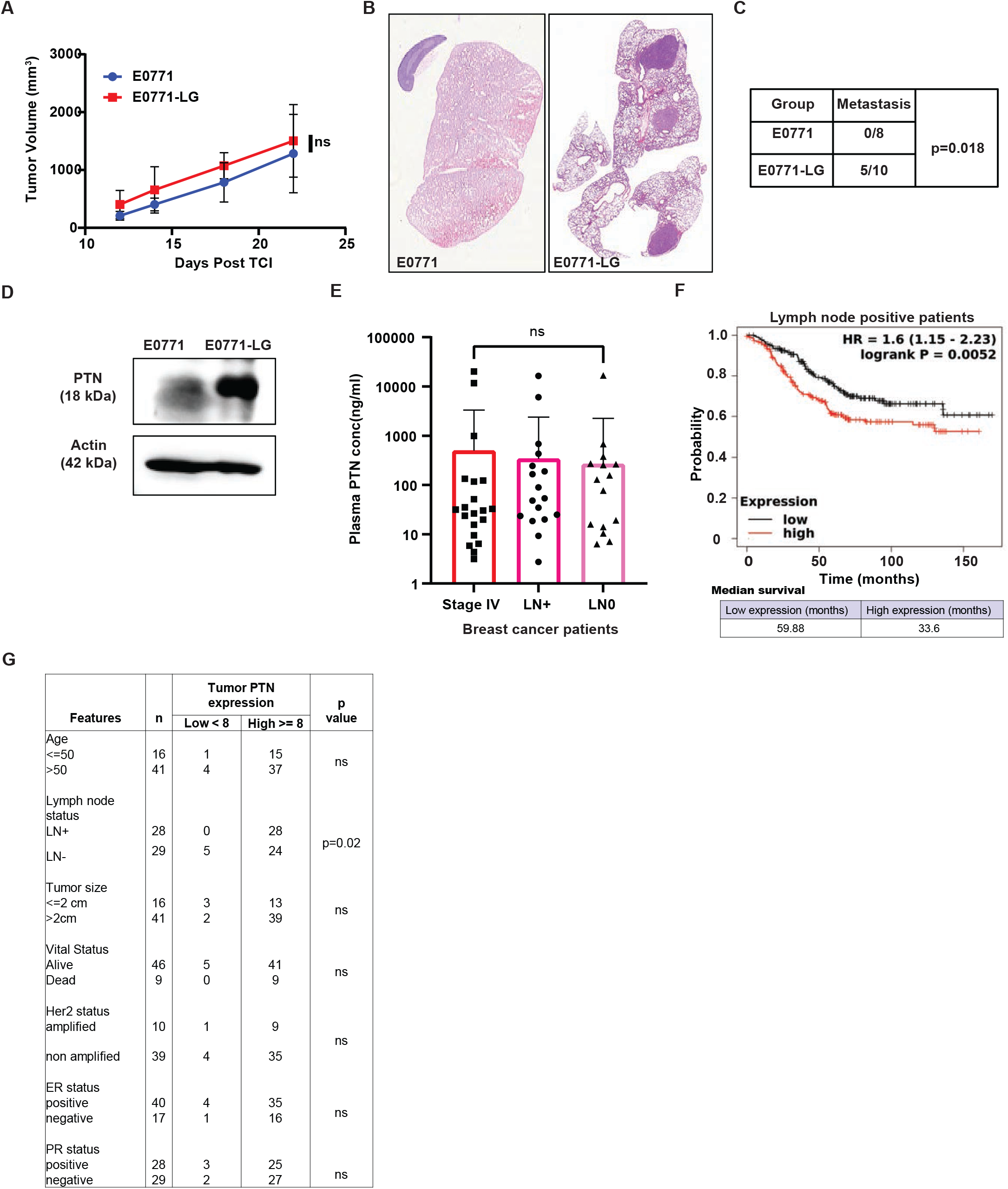
PTN is associated with metastasis in breast cancer. **A)** Growth curve of orthotopically implanted (4^th^ mammary fat pad) E0771 parental and E0771-LG tumors in wildtype (WT) C57BL/6 mice. **B)** Representative H&E images of lung metastasis from mice bearing E0771 or E0771-LG tumors. **C)** Quantification of gross metastasis to lungs in mice bearing E0771 or E0771-LG tumors. Chi-square test, *p<0.05. **D)** Western blot analysis of PTN expression (18kDa) in E0771 and E0771-LG cell lines, representative of three independent replicates. **E)** Bar graph showing plasma levels of PTN in stage IV (n=67), lymph node positive (n=72) and lymph node negative (n=71) breast cancer patients, as determined by ELISA. Statistical significance was determined using Kruskal-Wallis test. **F)** Kaplan Meier plot derived from TCGA database showing overall survival of lymph node positive breast cancer patients expressing high (n = 220) versus low (n = 232) levels of PTN in their primary tumor. Data were generated from https://kmplot.com/analysis/. **G)** Primary tumor tissues from Figure 2F were scored by a pathologist into PTN high and low expressing tumors. Additional patient history was obtained to determine correlation between PTN expression and clinical pathophysiological features of patients. Chi-square test, *p<0.05.

**Supplemental Figure 2.**
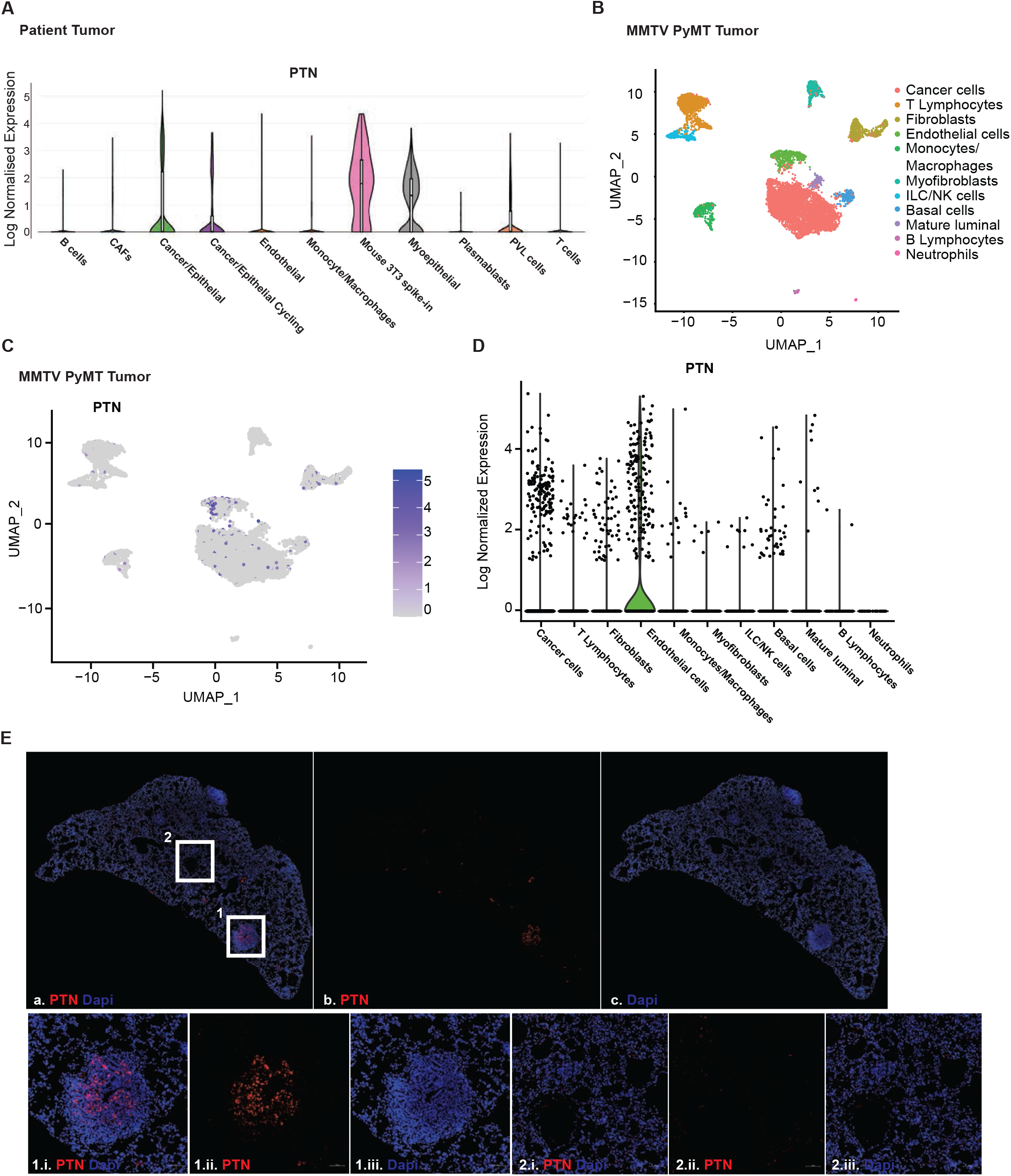
Pleiotrophin expression within different cell types of TME. **A)** Violin plot showing PTN expression in different cell types within the TME was generated using breast cancer patient scRNAseq data (BC-P1 CID4471) from a previous study (Wu et al., 2021) **B)** tSNE plot showing the different cell types identified from scRNAseq of late stage PyMT tumors. **C)** tSNE plot showing the expression of PTN in different cells from (B). **D)** Violin plot, generated from scRNAseq of MMTV-PyMT primary tumors, showing PTN expression by different cell types within the TME. **E)** Representative image of RNA FISH of PTN (red) on a lung bearing metastatic lesion from a MMTV-PyMT mice. Boxes 1 and 2 indicate metastatic lesion and normal lung area, respectively. Magnified images at 10x of metastatic lesion and normal lung area are shown in 1,i, 1,ii, 1,iii and 2.i, 2.ii, 2.iii, respectively. Scale bar, 50 μm.

**Supplemental Figure 3.**
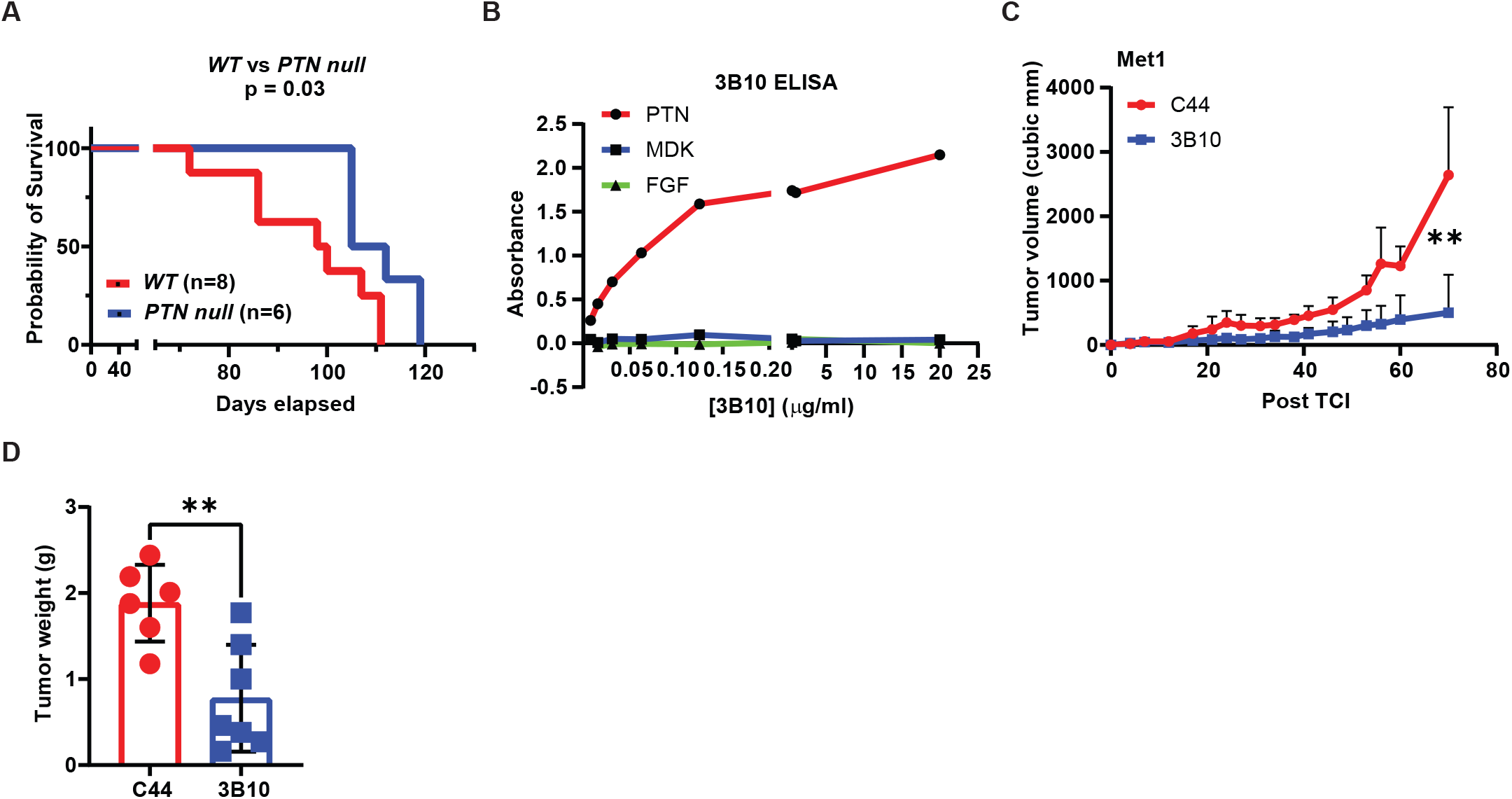
In vivo inhibition of PTN using a mouse monoclonal antibody, 3B10. **A)** Kaplan Meier plot comparing the overall survival of MMTV PyMT *WT* versus *PTN null* mice. Log-rank (Mantel-Cox) test, *p<0.05. **B)** Specificity of 3B10 for PTN in ELISA. Midkine (MDK) and Fibroblast growth factor (FGF) were used as negative controls. **C)** Growth curve of orthotopically implanted Met-1 tumors (MMTV-PyMT derived) in *WT* FVB mice treated with IgG control (C44) or anti-PTN therapy (3B10). Treatment started once tumors were palpable (~50 mm^3^). **D)** Met-1 tumor weight at the end of study in mice. Unpaired t-test, **p<0.01.

**Supplemental Figure 4.**
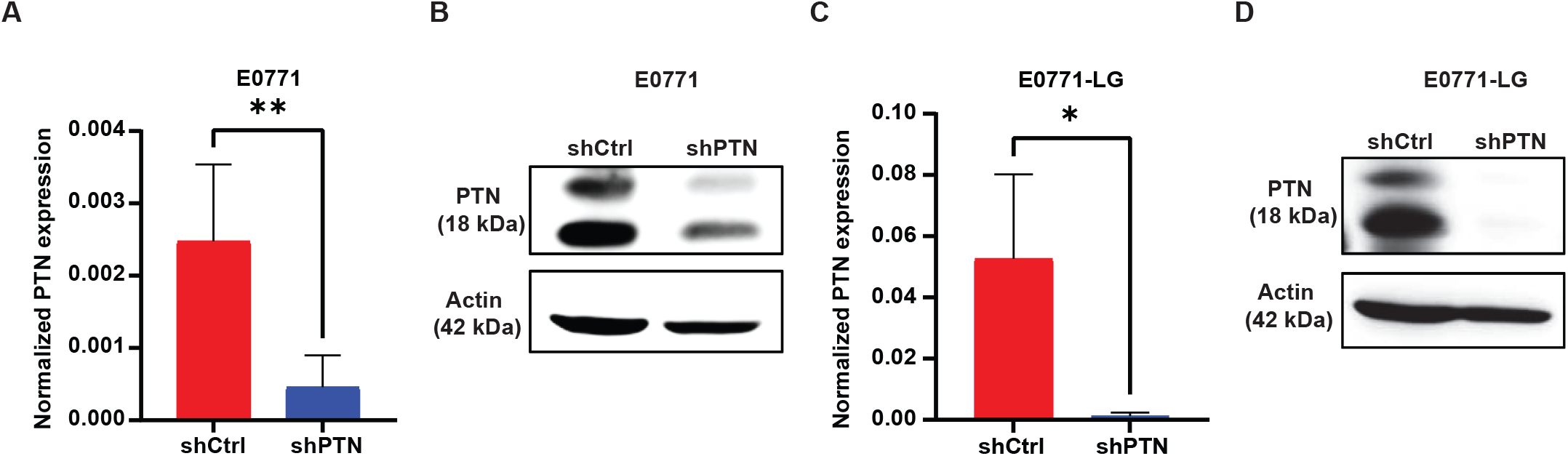
Validation of knockdown of PTN expression in E0771 and E0771-LG cells. **A)** Normalized PTN expression as determined by qPCR in E0771shCtrl and PTN knockdown cells, E0771shPTN. Unpaired t-test, **p<0.01. **B)** Western blot analysis of PTN expression (18kDa) in E0771shCtrl and E0771shPTN cell lines, representative of three independent replicates. **C)** Normalized PTN expression as determined by qPCR in E0771-LGshCtrl and PTN knockdown cells, E0771-LGshPTN. Unpaired t-test, *p<0.01. **D)** Western blot analysis of PTN expression (18kDa) in E0771-LGshCtrl and E0771-LGshPTN cell lines, representative of three independent replicates.

**Supplemental Figure 5.**
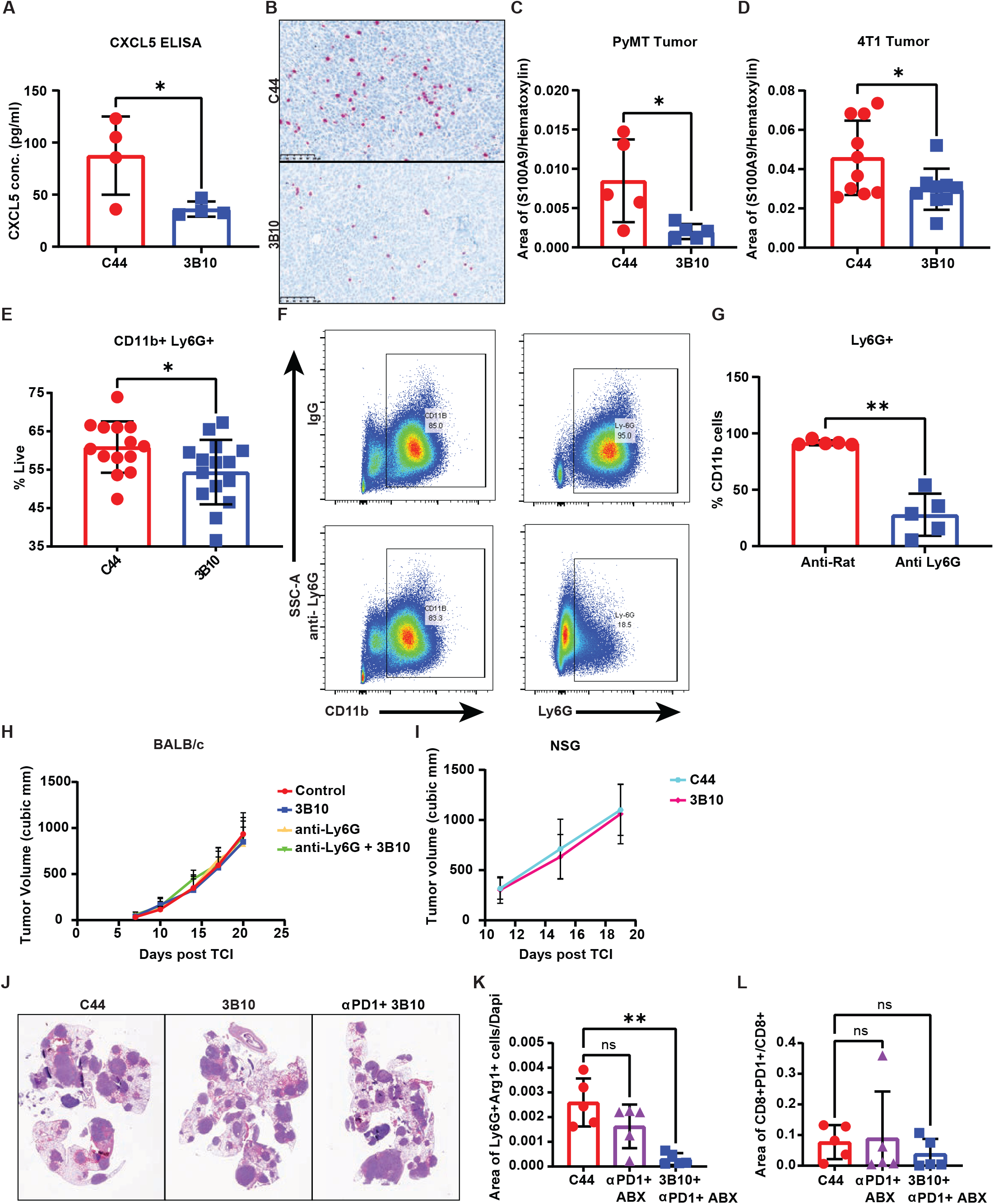
Effects of PTN inhibition on tumor phenotype. **A)** CXCL5 expression MMTV-PyMT tumors treated with C44 or 3B10 was determined by ELISA. Unpaired t-test, *p<0.05. **B)** 4T1 and MMTV-PyMT tumors from mice treated with C44 or 3B10 were stained for MDSCs (S100A9+ cells) by IHC. Representative images are shown at 20x magnification. Scale bar, 100 μm. **C-D)** Quantification of B. Unpaired t-test, *p<0.05. **E)** Quantification of neutrophils (CD11b+ Ly6G+) in 4T1 metastatic lungs from mice treated with C44 or 3B10 as determined by flow cytometry. Unpaired t-test, *p<0.05. **F)** Confirmation of Ly6G depletion by flow cytometry analysis of CD11b+ Ly6G+ cells in blood from mice bearing 4T1 tumor at the end of study. **G)** Quantification of F. Unpaired t-test, **p<0.01. **H)** Growth curve of orthotopically implanted 4T1 tumors in wildtype (WT) *BALB/c* mice treated with IgG control, 3B10, Ly6G or Ly6G+3B10. **I)** Growth curve of orthotopically implanted 4T1 tumors in NSG mice treated with C44 or 3B10. **J)** Representative H&E images of metastatic lungs from experimental metastasis study using E0771-LG cells in WT C57BL/6 mice that were treated with C44, 3B10 or anti-PD1+3B10. Treatment started on day 7 after tumor cell injection. **K-L)** Metastatic lungs from Figure 7G were stained by IHC and quantified for Ly6G+Arg1+ cells (**K**) and PD1+CD8+ cells (**L**). Kruskal-Wallis test, **p<0.01.

## Methods

### Cells

Murine triple negative breast adenocarcinoma (TNBC) cell line, 4T1 (CRL-2539) and immortalized human embryonic kidney cell line, HEK293T (CRL-3216), were obtained from ATCC. MMTV-PyMT (FVB/N) mammary adenocarcinoma derived cell line, Met-1(Borowsky et al., 2005), was kindly provided by Dr. Philip Thorpe (UT Southwestern Medical Center). Syngeneic murine TNBC(Johnstone et al., 2015) cell line, E0771 (CRL-3461) and E0771-LG(Kitamura et al., 2019), were gifts from Dr. Thorpe (UT Southwestern Medical Center) and Dr. Jeffrey Pollard (University of Edinburgh), respectively. Cells, PyMT and PyMT-LG, were derived from mammary adenocarcinoma and metastatic lungs of a 13-week-old MMTV-PyMT FVB/N female mice, respectively. For this, tumors and lungs were digested with a cocktail containing collagenase I (45 μ/ml; Worthington), collagenase II (15 μ/ml; Worthington), collagenase III (45 μ/ml; Worthington), collagenase IV (45 μ/ml; Worthington), elastase (0.075 μ/ml; Worthington), hyaluronidase (30 μ/ml; MilliporeSigma), and DNase type I (25 μ/ml; MilliporeSigma) for 1 hour at 37°C and passed through a 100 μm cell strainer (Falcon). PyMT and PyMT-LG cells were passaged at least 10 times before use. Mouse hybridoma producing mAb 3B10 (patent number: US 2014/0294845A1) specific for PTN was grown and supernatant was used to purify 3B10 through standard protein A chromatography. All the above-mentioned cells were cultured in DMEM (Invitrogen) containing 10% FBS and 1% v/v penicillin/streptomycin (Fisher Scientific) at 37°C in a humidified incubator with 5% CO_2_. Cells were confirmed to be free of pathogen before use.

### Animal studies

MMTV-PyMT FVB/N, FVB/N, NSG, BALB/c and C57BL/6 mice were purchased from The Jackson Laboratory. NSG breeder pairs were bred in-house at UT Southwestern. *Ptn-null* C57BL/6(Muramatsu et al., 2006) mice colony was provided by Dr. John Chute (Cedars Sinai). *Ptn-null* C57BL/6 mice were crossed with WT FVB/N mice to generate *Ptn-null* mice in FVB/N background. These were further crossed with MMTV-PyMT FVB/N males to develop MMTV-PyMT *Ptn-null* FVB/N mice. Newly generated mouse colonies were bred for 6 generations at minimum before use. For in vivo tumor studies, mouse breast cancer cells, 4T1 (1 × 10^5^), E0771 (5 × 10^5^), E0771-LG (5 × 10^5^) and Met-1 (5 × 10^5^) were injected orthotopically into the 4^th^ mammary fat pad of 8-week-old female BALB/c, C57BL/6 or FVB/N mice. Tumors were measured twice weekly and tumor volumes were calculated using the formula: V = (a × b^2^)/2, where a is length and b is width. Treatments started when tumors were palpable (~50 mm^3^). Mice were euthanized when tumor volume from the control group reached 2000 mm^3^. For survival studies, mice were euthanized when tumors reached maximum allowable limit. For experimental metastasis assay, E0771-LG (5 × 10^5^) cells were injected via the tail vein and mice were euthanized on day 19. Endpoint of experimental metastasis assay was selected based on pilot studies (data not shown). For MMTV-PyMT therapy experiments, treatment started at 8 weeks of age. For all in vivo studies, mice were randomized into different treatment groups. Please refer to the figure legend of each figure for exact therapy window and treatment groups. Treatment dosage consisted of mouse IgG control C44 (15 mg/kg, CRL-1943, twice per week), 3B10 (15 mg/kg, twice per week), anti-PD-1 (5 mg/kg, #BE0273, BioXCell, twice per week), abraxane (25 mg/kg, #68817-134-50, Celgene, twice per week), anti-Rat IgG control (10 mg/kg, #BP0089, BioXCell, every three days) and anti-Ly6G (10 mg/kg, BP0075-1, BioXCell, every three days). For endpoint experiments all mice were sacrificed at the same time. Tumors and lungs were weighed and gross metastasis to lungs were counted. Tissues were fixed in 10% formalin for 48 hours or snap frozen in liquid nitrogen for further studies or digested into single-cell suspension for flow cytometry. All animals were housed in a pathogen-free facility with 24-hour access to food and water ad libitum. All animal procedures were approved and in compliance with IACUC policies.

### Clinical sample procurement and inclusion criteria

Lymph node and tumor FFPE sections as well as blood plasma from breast cancer patients were obtained from UTSW tissue management shared resources. Three cohorts of patient samples were included in this study, namely, stage IV, lymph node positive and lymph node negative patients. Inclusion criteria for cohort selection comprised of patients with stage IV disease, patients with lymph node metastases of at least 2 mm or larger and patients with microscopically negative lymph node metastasis, respectively. Case details including overall survival, time of diagnosis, cancer subtype etc. were also obtained for each patient.

### ELISA and cytokine array

Patient blood samples were collected in EDTA tubes. Plasma samples were procured from UTSW tissue management shared resources. These plasma samples were diluted at least 2x or more (as required) and then were used to detect PTN by ELISA using commercial human PTN ELISA kits (ThermoFisher Scientific, EH370RB). CXCL5 ELISA from tumor lysates was performed using commercial kit (R&D Systems, MX000) by following their instructions manual.

Mouse cytokine antibody array – abeam (ab133994) and RayBiotech (C3AAM-CYT-3-2) were used to detect cytokines from mouse PyMT derived tumor (Met-1) and 4T1 lung lysates, respectively.

### Survival curve generation

Patients were segregated into ‘PTN high’ or ‘PTN low’ groups based on plasma PTN concentration. PTN concentrations above detectable level, 1 ng/ml, were considered ‘PTN high’ while those lower were considered ‘PTN low’. Overall survival data, obtained from UTSW tissue management shared resources, were plotted for ‘PTN high’ versus ‘PTN low’ cohort of patients in Graphpad Prism9. Log rank (Mantel-Cox) test was used to determine significance and hazard ratio. Kaplan Meier plot for overall survival of stage IV and lymph node positive breast cancer patients based on PTN RNA expression were generated from https://kmplot.com/analvsis/.

### Immunohistochemistry (IHC)

IHC was performed using previously described (Sorrelle et al., 2019) protocol. Briefly, slides were warmed in a 60°C oven for 10 min before deparaffinization and rehydration. Slides were fixed in 10% neutral buffered formalin for 30 min followed by a PBS wash. Antigen retrieval was performed in antigen retrieval buffer (10 mM Tris-HCI, 1 mM EDTA with 10% glycerol) at 110°C for 18 min (~4-5ψ). Tissue sections were blocked with 2.5% horse serum (Vector Laboratories, S-2012-50) or 2.5% goat serum (Vector Laboratories, S-1012-50) followed by overnight incubation with primary antibody. Next day, slides were washed and incubated with HRP conjugated secondary antibody for 30 min on a shaker. Slides were then washed three times for 5 min in PBST before developing signal. For developing chromogen signal, Bentazoid DAB (BDB2004L) was used. Slides were counter-stained with hematoxylin and then cover-slipped using VectaMount (H-5501, Vector Laboratories) and scanned at 20X using the Hamamatsu NanoZoomer 2.0-HT. For developing fluorescence signal, Opal substrates (Akoya Biosciences, SKUFP1488001KT, SKU FP1487001KT, SKUFP1497001KT) were used. When performing multiplex IHC, slides were stripped off of antibodies by incubating in 10 mM citrate buffer (pH 6.2, 10% glycerol) in a pressure cooker at 110°C for 2 min. Sequential staining was performed by stripping after every round of staining before probing for the next antibody. Slides were counter-stained with DAPI and then cover-slipped using Prolong Gold (#P36931, Life Technologies). Slides were scanned at 40X using the Zeiss Axioscan.Z1 in Whole Brain Microscopy Facility of UT Southwestern.

### Scoring of clinical slides by pathologist

Patient tumor tissues stained for PTN by IHC were scored by a pathologist Dr. Yisheng Fang. Briefly, PTN expression were evaluated by the proportion and intensity of the stained cells. The percentage of the stained tumor cells was categorized as follows: 0 (<5%), 1 (5%—25%), 2 (26%-50%), 3 (51%-75%) and 4 (>75%).The cytoplasmic staining intensity was classified as: 0 (negative), 1 (weak, defined as barely discernable at 4X, visible at 10X), 2 (moderate, defined as visible at 4X) and 3 (strong, dark brown obvious at 4X). The final score was obtained by multiplication of the percentage of stained cells and staining intensity to obtain a total score ranging from 0 to 12. Score of less than 8 was considered ‘PTN low’ while the rest was considered ‘PTN high’.

### Antibodies and reagents

For IHC, the following primary antibodies were used: anti-human PTN (1:200; R&D systems, AF252-PB), anti-PanCK, (1:500, Novus Biologicals, NBP2-29429); anti-human CD31 (1:500, Invitrogen, 14-0319-82), anti-PyMT (1:200, Santa Cruz, sc53481), antimouse CD31 (1:500, cell signaling 77699S), anti-Podoplanin (1:200, AB11936, abeam), anti-CD45 (1:500, Cell Signaling, 70257), anti-mouse Ly-6G (1:200, BD Biosciences,551459), anti-mouse S100A9 (1:500, Cell Signaling,73425), anti-CD3e (1:500, ThermoFisher,PA1-29547), anti-mouse CD8 (1:1000, Cell Signaling, 98941S), anti-iNos (1:200, ThermoFisher, PA1-21054), anti-Arg1(1:500, Cell Signaling, 93668S), anti-Granzyme B (1:500, Cell Signaling, 46890S). Secondary antibodies used for IHC included: HRP conjugated secondary anti-goat antibody (ImmPRESS; Vector Laboratories, MP-7405-50) or anti-rabbit antibody (ImmPRESS; Vector Laboratories, MP-7451) or anti-rat antibody (ImmPRESS; Vector Laboratories, MP-7404-50) or HRP anti-syrian hamster antibody (1:500, Jackson ImmunoResearch, 107-035-142).

For immunoblotting the following primary antibodies were used: anti-PTN (1:1000; R&D systems, AF252-PB), anti-p65 (1:1000, Cell Signaling, 8242), anti-p-p65 (1:1000, Cell Signaling, 3033S), anti-actin (1:2500, Sigma-Aldrich, A2066). Anti-goat HRP (1:10,000, Jackson ImmunoResearch, HRP 705-035-147) and anti-rabbit HRP (1:10,000, Jackson ImmunoResearch, 711-035-152) were used as secondary antibodies corresponding to the host of primary. Recombinant mouse PTN (6580-PL) was used for in vitro cancer cell stimulation experiments.

### Quantification of Immunohistochemistry

Quantification of IHC was done using Fiji software. DAB intensity was quantified as previously described(Nguyen DH, 2013). Briefly, representative images were taken from each section. Each image was color deconvoluted into DAB and hematoxylin. Five random areas were selected to measure mean gray value. Average mean gray value (f) was obtained, and intensity was calculated using the formula: Intensity = 250 - f. To calculate positively stained area, the deconvoluted DAB image was threshold at the same intensity value for PTN staining and area of threshold was measured. All calculated areas were normalized using hematoxylin. Quantification represents PTN positive area/hematoxylin area. For fluorescent images, composite images were created. Single color and co-localized area were calculated by thresholding for each color. All area calculations were normalized to Dapi.

### Single Cell RNA sequencing (scRNAseq) data collection

Breast cancer patient scRNAseq data (BC-P1 CID4471) was accessed from Wu et al(Wu et al., 2021) using the following online portal: https://sinalecell.broadinstitute.org/. UMAPs and violin plot of PTN expression were generated using BC-P1 CID4471 patient data from this study. Mouse MMTV-PyMT mammary tumor scRNAseq data was accessed from a previous study(Valdes-Mora et al., 2021). UMAPs and violin plot of PTN expression were generated from MMTV PyMT/WT data set.

### Dual Immunohistochemistry (IHC) and Fluorescence in situ hybridization (FISH)

FISH on FFPE tissue or fixed cultured cells was performed following the instructions in RNAscope kit (catalogue number: 323110, advanced cell diagnostics INC). Ptn (catalogue number: 451181), RPTPβ/ζ or PTPRZ1 (#460991), positive control (catalogue number: 313911), and negative control (catalogue number: 310043) probes were purchased from advanced cell diagnostics INC. When performing dual FISH and IF, following FISH, HRP blocker (provided in the kit) was used to block further HRP activity. From here on, slides were stained by IHC as detailed in the IHC method section above. For stripping process, slides were incubated in 10 mM citrate buffer (pH 6.2, 10% glycerol) in a pressure cooker at 110°C for 2 min before probing with the next primary antibody. Slides were counter-stained with DAPI and then cover-slipped using Prolong Gold (#P36931, Life Technologies). Slides were scanned at 40X using the Zeiss Axioscan.Z1 in Whole Brain Microscopy Facility of UT Southwestern. FISH on fixed cells were scanned at 40x using a laser scanning confocal microscope Zeiss LSM780 at Live Cell Imaging core of UT Southwestern.

### Transfection and knock-down

HEK293T cells were transfected with Lipofectamine 2000 reagent (Thermofisher,11668027) at 70% confluency. The shRNA construct used was PTN KD (TRCN0000071677, Sigma). For a negative control, pLKO.1-puro carrying a sequence targeting EGFP (Sigma) was used. E0771 and E0771-LG target cells were infected with virus thrice before selection under 3ug/ml of puromycin. Cells were maintained in puromycin for 6 weeks before use.

### Immunoblotting

Cell were lysed in ice cold RIPA buffer (50 mM Tris—HCI, pH 8.0, 150 mM NaCI, 0.1% SDS, 0.5% sodium deoxycholate, and 1% Nonidet P-40) for SDS-PAGE. Protein concentration was determined using a BCA Protein Assay Kit (23225, Thermo Fisher Scientific). Samples were boiled for 5 min in laemmli sample buffer. The proteins were then resolved by SDS-PAGE, transferred to nitrocellulose membranes, and blocked in WestVision (SP-7000-500) block. Membranes were incubated with primary antibody overnight. Next day, membranes were washed thrice for 5 min with TBST, and incubated with horseradish peroxidase-conjugated secondary antibody (Jackson ImmunoResearch Laboratories) for 1 h. Excess secondary antibody was thoroughly removed by washing 3 × 10min with TBST. The membranes were developed with SuperSignal West Pico substrate (34580, Thermo Fisher Scientific). Licor Odyssey Fc imaging system was used for the detection of the immunoreactive bands.

### Flow cytometry

Lungs bearing metastasis were digested with a cocktail containing collagenase I (45 U/mL; Worthington), collagenase II (15 U/mL; Worthington), collagenase III (45 U/mL; Worthington), collagenase IV (45 U/mL; Worthington), elastase (0.075 U/mL; Worthington), hyaluronidase (30 U/mL; MilliporeSigma), and DNase type I (25 U/mL; MilliporeSigma) for 60 min at 37°C and passed through a 70 μm cell strainer (Falcon). Whole blood collected from tumor bearing mice was spun 2000 rpm for 20 min at 4°C to collect cell pellet. Cells from blood and lungs were washed with PBS and stained with Ghost Viability Dye 510 (BD Biosciences) for 15 min. Cells were then washed and stained with antibodies against CD11b (BD Biosciences, 557657) and Ly-6G (BD Biosciences, 740953) for 1 hour at 4°C. Cells were analyzed using FACS LSRFortessa SORP (BD Biosciences) with the help of the Moody Foundation flow cytometry facility at UT Southwestern Medical Center. Analysis was performed using FlowJo software.

### Tissue histology

Tissues were fixed in 10% neutral buffered formalin for 48 hours, embedded in paraffin blocks and cut into 5 μm sections. H&E was performed, and slides were scanned using at 20X using Hamamatsu NanoZoomer 2.0-HT at Whole Brain Microscopy Facility of UT Southwestern. Lung H&Es were evaluated for metastasis. Metastasis was quantitated as area of metastasis/ total lung area. Areas were determined using NDPview software.

### RNA isolation and qPCR

RNA was isolated from cells and frozen tumor tissues in RLT buffer, and the protocol provided in QIAGEN RNAeasy plus mini kit (#74034). RNA concentration and purity were measured using a NanoDrop spectrophotometer. cDNAs were synthesized from 1 μg of RNA using iScript cDNA synthesis kit (Bio-rad, #1708891). 25 ng of total cDNA was used for qPCR with iTaq ™ Universal SYBR^®^ Green Supermix (Biorad, #172-5121). Samples were run in three technical replicates on a CFX96 real time system (Bio-rad). Assays consisted of three biological replicates. Primers used were as follow: msPTN (fwd: 5’CTCTGCACAATGCTGACTGTC3’; rev: 5’CTTTGACTCCGCTTGAGGCTT3’).

### mRNA sequencing and downstream analysis

Total RNA was isolated from tumors and cell lines from three biological replicates and shipped to Novogene Inc. (Sacramento, CA). RNA integrity scores (RIN score) for samples were greater than 8. Sequencing was performed on an lIumina hiseq4000 sequencer (paired end 150nt) resulting in an average of 43 million reads of which an average of 93% mapping to the mm10 reference genome. Fastq files were quality controlled with fastqc (https://www.bioinformatics.babraham.ac.uk/projects/fastqc/). Then aligned assembled with the Hisat2/Stringtie pipeline (http://daehwankimlab.github.io/hisat2/, https://ccb.jhu.edu/software/stringtie/)(Kim et al., 2019). Differential analysis was performed in R using Rsubread, (https://bioconductor.org/packages/release/bioc/html/Rsubread.html (Liao et al., 2019) and edgeR https://bioconductor.org/packages/release/bioc/html/edgeR.html)(Robinson et al., 2010). Statistical analysis was used using the robust quasi-likelihood pipeline by edgeR. Metascape was used to perform Motif enrichment and TRRUST analysis on differentially expressed genes(Han et al., 2018). CIBERSORT analysis was performed at TIMER2.0 using immune estimation feature. Heat map of neutrophil associated genes was generated in Graphpad prism.

### Statistics

All analysis were performed in Graphpad prism 9. All mice used in this manuscript were females of similar age. All data is represented as mean ± SD. Two tailed student’s t test (paired or unpaired) and one-way ANOVA were used to compare two or more groups respectively. All experiments were performed in at least three biological replicates. A p value of less than 0.05 was considered significant.

## Data availability

All raw data generated in this manuscript has been deposited in Gene Expression Omnibus (accession number will be provided upon revision).

## Schematic

All schematic cartoons were made using Biorender.

## References

Albrengues, J., M.A. Shields, D. Ng, C.G. Park, A. Ambrico, M.E. Poindexter, P. Upadhyay, D.L. Uyeminami, A. Pommier, V. Kuttner, E. Bruzas, L. Maiorino, C. Bautista, E.M. Carmona, P.A. Gimotty, D.T. Fearon, K. Chang, S.K. Lyons, K.E. Pinkerton, L.C. Trotman, M.S. Goldberg, J.T. Yeh, and M. Egeblad. 2018. Neutrophil extracellular traps produced during inflammation awaken dormant cancer cells in mice. Science 361:

Attalla, S., T. Taifour, T. Bui, and W. Muller. 2021. Insights from transgenic mouse models of PyMT-induced breast cancer: recapitulating human breast cancer progression in vivo. Oncogene 40:475–491.

Averill, M.M., S. Barnhart, L. Becker, X. Li, J.W. Heinecke, R.C. Leboeuf, J.A. Hamerman, C. Sorg, C. Kerkhoff, and K.E. Bornfeldt. 2011. S100A9differentially modifies phenotypic states of neutrophils, macrophages, and dendritic cells: implications for atherosclerosis and adipose tissue inflammation. Circulation 123:1216–1226.

Ben-Porath, I., M.W. Thomson, V.J. Carey, R. Ge, G.W. Bell, A. Regev, and R.A. Weinberg. 2008. An embryonic stem cell-like gene expression signature in poorly differentiated aggressive human tumors. Nat Genet 40:499–507.

Borowsky, A.D., R. Namba, L.J. Young, K.W. Hunter, J.G. Hodgson, C.G. Tepper, E.T. McGoldrick, W.J. Muller, R.D. Cardiff, and J.P. Gregg. 2005. Syngeneic mouse mammary carcinoma cell lines: two closely related cell lines with divergent metastatic behavior. Clin Exp Metastasis 22:47–59.

Cameron, M.D., E.E. Schmidt, N. Kerkvliet, K.V. Nadkarni, V.L. Morris, A.C. Groom, A.F. Chambers, and I.C. MacDonald. 2000. Temporal progression of metastasis in lung: cell survival, dormancy, and location dependence of metastatic inefficiency. Cancer Res 60:2541–2546.

Celia-Terrassa, T., and Y. Kang. 2016. Distinctive properties of metastasis-initiating cells. Genes Dev 30:892–908.

Cerezo-Wallis, D., M. Contreras-Alcalde, K. Troule, X. Catena, C. Mucientes, T.G. Calvo, E. Canon, C. Tejedo, P.C. Pennacchi, S. Hogan, P. Kolblinger, H. Tejero, A.X. Chen, N. Ibarz, O. Grana-Castro, L. Martinez, J. Munoz, P. Ortiz-Romero, J.L. Rodriguez-Peralto, G. Gomez-Lopez, F. Al-Shahrour, R. Rabadan, M.P. Levesque, D. Olmeda, and M.S. Soengas. 2020. Midkine rewires the melanoma microenvironment toward a tolerogenic and immune-resistant state. Nat Med 26:1865–1877.

Chambers, A.F., A.C. Groom, and I.C. MacDonald. 2002. Dissemination and growth of cancer cells in metastatic sites. Nat Rev Cancer 2:563–572.

Chang, Y., M. Zuka, P. Perez-Pinera, A. Astudillo, J. Mortimer, J.R. Berenson, and T.F. Deuel. 2007. Secretion of pleiotrophin stimulates breast cancer progression through remodeling of the tumor microenvironment. Proc Natl Acad Sci U S A 104:10888–10893.

Chao, T., E.E. Furth, and R.H. Vonderheide. 2016. CXCR2-DependentAccumulation of Tumor-Associated Neutrophils Regulates T-cell Immunity in Pancreatic Ductal Adenocarcinoma. Cancer Immunol Res 4:968–982.

Deng, J., Y. Kang, C.C. Cheng, X. Li, B. Dai, M.H. Katz, T. Men, M.P. Kim, E.A. Koay, H. Huang, R.A. Brekken, and J.B. Fleming. 2021. DDR1-induced neutrophil extracellular traps drive pancreatic cancer metastasis. JCI Insight 6:

Ewens, A., E. Mihich, and M.J. Ehrke. 2005. Distant metastasis from subcutaneously grown E0771 medullary breast adenocarcinoma. Anticancer Res 25:3905–3915.

Garver, R.I., Jr., D.M. Radford, H. Donis-Keller, M.R. Wick, and P.G. Milner. 1994. Midkine and pleiotrophin expression in normal and malignant breast tissue. Cancer 74:1584–1590.

Giamanco, N.M., Y.H. Jee, A. Wellstein, C.D. Shriver, T.A. Summers, and J. Baron. 2017. Midkine and pleiotrophin concentrations in needle biopsies of breast and lung masses. Cancer Biomark 20:299–307.

Gu, D., B. Yu, C. Zhao, W. Ye, Q. Lv, Z. Hua, J. Ma, and Y. Zhang. 2007. The effect of pleiotrophin signaling on adipogenesis. FEBS Lett 581:382–388.

Gutmann, D.H. 2017. TheTropism of Pleiotrophin: Orchestrating Glioma Brain Invasion. Cell 170:821–822.

Han, H., J.W. Cho, S. Lee, A. Yun, H. Kim, D. Bae, S. Yang, C.Y. Kim, M. Lee, E. Kim, S. Lee, B. Kang, D. Jeong, Y. Kim, H.N. Jeon, H. Jung, S. Nam, M. Chung, J.H. Kim, and I. Lee. 2018. TRRUST v2: an expanded reference database of human and mouse transcriptional regulatory interactions. Nucleic Acids Res 46:D38O–D386.

Himburg, H.A., G.G. Muramoto, P. Daher, S.K. Meadows, J.L. Russell, P. Doan, J.T. Chi, A.B. Salter, W.E. Lento, T. Reya, N.J. Chao, and J.P. Chute. 2010. Pleiotrophin regulates the expansion and regeneration of hematopoietic stem cells. Nat Med 16:475–482.

Huang, P., D.J. Ouyang, S. Chang, M.Y. Li, L. Li, Q.Y. Li, R. Zeng, Q.Y. Zou, J. Su, P. Zhao, L. Pei, and W.J. Yi. 2018. Chemotherapy-driven increases in the CDKN1A/PTN/PTPRZ1 axis promote chemoresistance by activating the NF-kappaB pathway in breast cancer cells. Cell Commun Signal 16:92.

Imai, S., M. Kaksonen, E. Raulo, T. Kinnunen, C. Fages, X. Meng, M. Lakso, and H. Rauvala. 1998. Osteoblast recruitment and bone formation enhanced by cell matrix-associated heparin-binding growth-associated molecule (HB-GAM). J Cell Biol 143:1113–1128.

Johnstone, C.N., Y.E. Smith, Y. Cao, A.D. Burrows, R.S. Cross, X. Ling, R.P. Redvers, J.P. Doherty, B.L. Eckhardt, A.L. Natoli, C.M. Restall, E. Lucas, H.B. Pearson, S. Deb, K.L. Britt, A. Rizzitelli, J. Li, J.H. Harmey, N. Pouliot, and R.L. Anderson. 2015. Functional and molecular characterisation of E0771.LMB tumours, a new C57BL/6-mouse-derived model of spontaneously metastatic mammary cancer. Dis Model Mech 8:237–251.

Kim, D., J.M. Paggi, C. Park, C. Bennett, and S.L. Salzberg. 2019. Graph-based genome alignment and genotyping with HISAT2 and HISAT-genotype. Nat Biotechnol 37:907–915.

Kitamura, T., Y. Kato, D. Brownlie, D.Y.H. Soong, G. Sugano, N. Kippen, J. Li, D. Doughty-Shenton, N. Carragher, and J.W. Pollard. 2019. Mammary Tumor Cells with High Metastatic Potential Are Hypersensitive to Macrophage-Derived HGF. Cancer Immunol Res 7:2052–2064.

Landgraf, P., P. Wahle, H.C. Pape, E.D. Gundelfinger, and M.R. Kreutz. 2008. The survival-promoting peptide Y-P30 enhances binding of pleiotrophin to syndecan-2 and-3 and supports its neuritogenic activity. J Biol Chem 283:25036–25045.

Lawson, D.A., N.R. Bhakta, K. Kessenbrock, K.D. Prummel, Y. Yu, K. Takai, A. Zhou, H. Eyob, S. Balakrishnan, C.Y. Wang, P. Yaswen, A. Goga, and Z. Werb. 2015. Single-cell analysis reveals a stem-cell program in human metastatic breast cancer cells. Nature 526:131–135.

Li, F., F. Tian, L. Wang, I.K. Williamson, B.G. Sharifi, and P.K. Shah. 2010. Pleiotrophin (PTN) is expressed in vascularized human atherosclerotic plaques: IFN-{gamma}/JAK/STAT1 signaling is critical for the expression of PTN in macrophages. FASEB J 24:810–822.

Li, Y.S., P.G. Milner, A.K. Chauhan, M.A. Watson, R.M. Hoffman, C.M. Kodner, J. Milbrandt, and T.F. Deuel. 1990. Cloning and expression of a developmentally regulated protein that induces mitogenic and neurite outgrowth activity. Science 250:1690–1694.

Liao, Y., G.K. Smyth, and W. Shi. 2019. The R package Rsubread is easier, faster, cheaper and better for alignment and quantification of RNA sequencing reads. Nucleic Acids Res 47:e47.

Lin, E.Y., J.G. Jones, P. Li, L. Zhu, K.D. Whitney, W.J. Muller, and J.W. Pollard. 2003. Progression to malignancy in the polyoma middle T oncoprotein mouse breast cancer model provides a reliable model for human diseases. Am J Pathol 163:2113–2126.

Liu, X., G.A. Mashour, H.F. Webster, and A. Kurtz. 1998. Basic FGF and FGF receptor 1 are expressed in microglia during experimental autoimmune encephalomyelitis: temporally distinct expression of midkine and pleiotrophin. Glia 24:390–397.

Luzzi, K.J., I.C. MacDonald, E.E. Schmidt, N. Kerkvliet, V.L. Morris, A.F. Chambers, and A.C. Groom. 1998. Multistep nature of metastatic inefficiency: dormancy of solitary cells after successful extravasation and limited survival of early micrometastases. Am J Pathol 153:865–873.

Ma, J., Y. Kong, H. Nan, S. Qu, X. Fu, L. Jiang, W. Wang, H. Guo, S. Zhao, J. He, and K. Nan. 2017. Pleiotrophin as a potential biomarker in breast cancer patients. Clin Chim Acta 466:6–12.

Muramatsu, H., P. Zou, N. Kurosawa, K. Ichihara-Tanaka, K. Maruyama, K. Inoh, T. Sakai, L. Chen, M. Sato, and T. Muramatsu. 2006. Female infertility in mice deficient in midkine and pleiotrophin, which form a distinct family of growth factors. Genes Cells 11:1405–1417.

Nguyen DH, Z.T., Shu J, and Mao JH. 2013. Quantifying chromogen intensity in immunohistochemistry via reciprocal intensity. Cancer InCytes

Olmeda, D., D. Cerezo-Wallis, E. Riveiro-Falkenbach, P.C. Pennacchi, M. Contreras-Alcalde, N. Ibarz, M. Cifdaloz, X. Catena, T.G. Calvo, E. Canon, D. Alonso-Curbelo, J. Suarez, L. Osterloh, O. Grana, F. Mulero, D. Megias, M. Canamero, J.L. Martinez-Torrecuadrada, C. Mondal, J. Di Martino, D. Lora, I. Martinez-Corral, J.J. Bravo-Cordero, J. Munoz, S. Puig, P. Ortiz-Romero, J.L. Rodriguez-Peralto, S. Ortega, and M.S. Soengas. 2017. Whole-body imaging of lymphovascular niches identifies pre-metastatic roles of midkine. Nature 546:676–680.

Papadimitriou, E., E. Pantazaka, P. Castana, T. Tsalios, A. Polyzos, and D. Beis. 2016. Pleiotrophin and its receptor protein tyrosine phosphatase beta/zeta as regulators of angiogenesis and cancer. Biochim Biophys Acta 1866:252–265.

Park, J., R.W. Wysocki, Z. Amoozgar, L. Maiorino, M.R. Fein, J. Jorns, A.F. Schott, Y. Kinugasa-Katayama, Y. Lee, N.H. Won, E.S. Nakasone, S.A. Hearn, V. Kuttner, J. Qiu, A.S. Almeida, N. Perurena, K. Kessenbrock, M.S. Goldberg, and M. Egeblad. 2016. Cancer cells induce metastasis-supporting neutrophil extracellular DNA traps. Sci Transl Med 8:361ra138.

Perez-Pinera, P., O. Garcia-Suarez, P. Menendez-Rodriguez, J. Mortimer, Y. Chang, A. Astudillo, and T.F. Deuel. 2007. The receptor protein tyrosine phosphatase (RPTP)beta/zeta is expressed in different subtypes of human breast cancer. Biochem Biophys Res Commun 362:5–10.

Pufe, T., M. Bartscher, W. Petersen, B. Tillmann, and R. Mentlein. 2003. Expression of pleiotrophin, an embryonic growth and differentiation factor, in rheumatoid arthritis. Arthritis Rheum 48:660–667.

Qin, E.Y., D.D. Cooper, K.L. Abbott, J. Lennon, S. Nagaraja, A. Mackay, C. Jones, H. Vogel, P.K. Jackson, and M. Monje. 2017. Neural Precursor-Derived Pleiotrophin Mediates Subventricular Zone Invasion by Glioma. Cell 170:845–859 e819.

Robinson, M.D., D.J. McCarthy, and G.K. Smyth. 2010. edgeR: a Bioconductor package for differential expression analysis of digital gene expression data. Bioinformatics 26:139–140.

Ryan, E., D. Shen, and X. Wang. 2016. Structural studies reveal an important role for the pleiotrophin C-terminus in mediating interactions with chondroitin sulfate. FEBS J 283:1488–1503.

Ryan, E., D. Shen, and X. Wang. 2021. Pleiotrophin interactswith glycosaminoglycans in a highly flexible and adaptable manner. FEBS Lett 595:925–941.

Sevillano, J., M.G. Sanchez-Alonso, B. Zapateria, M. Calderon, M. Alcala, M. Limones, J. Pita, E. Gramage, M. Vicente-Rodriguez, D. Horrillo, G. Medina-Gomez, M.J. Obregon, M. Viana, I. Valladolid-Acebes, G. Herradon, and M.P. Ramos-Alvarez. 2019. Pleiotrophin deletion alters glucose homeostasis, energy metabolism and brown fat thermogenicfunction in mice. Diabetologia 62:123–135.

Shi, Y., Y.F. Ping, W. Zhou, Z.C. He, C. Chen, B.S. Bian, L. Zhang, L. Chen, X. Lan, X.C. Zhang, K. Zhou, Q. Liu, H. Long, T.W. Fu, X.N. Zhang, M.F. Cao, Z. Huang, X. Fang, X. Wang, H. Feng, X.H. Yao, S.C. Yu, Y.H. Cui, X. Zhang, J.N. Rich, S. Bao, and X.W. Bian. 2017. Tumour-associated macrophages secrete pleiotrophin to promote PTPRZ1 signalling in glioblastoma stem cells for tumour growth. Nat Commun 8:15080.

Sorrelle, N., A.T.A. Dominguez, and R.A. Brekken. 2017. From top to bottom: midkine and pleiotrophin as emerging players in immune regulation. J Leukoc Biol 102:277–286.

Sorrelle, N., D. Ganguly, A.T.A. Dominguez, Y. Zhang, H. Huang, L.N. Dahal, N. Burton, A. Ziemys, and R.A. Brekken. 2019. Improved Multiplex Immunohistochemistry for Immune Microenvironment Evaluation of Mouse Formalin-Fixed, Paraffin-Embedded Tissues. J Immunol 202:292–299.

Sugiura, K., and C.C. Stock. 1952. Studies in a tumor spectrum. I. Comparison of the action of methylbis (2-chloroethyl)amine and 3-bis(2-chloroethyl)aminomethyl-4-methoxymethyl-5-hydroxy-6-methylpyridine on the growth of a variety of mouse and rat tumors. Cancer 5:382–402.

Szczerba, B.M., F. Castro-Giner, M. Vetter, I. Krol, S. Gkountela, J. Landin, M.C. Scheidmann, C. Donato, R. Scherrer, J. Singer, C. Beisel, C. Kurzeder, V. Heinzelmann-Schwarz, C. Rochlitz, W.P. Weber, N. Beerenwinkel, and N. Aceto. 2019. Neutrophils escort circulating tumour cells to enable cell cycle progression. Nature 566:553–557.

Valdes-Mora, F., R. Salomon, B.S. Gloss, A.M.K. Law, J. Venhuizen, L. Castillo, K.J. Murphy, A. Magenau, M. Papanicolaou, L. Rodriguez de la Fuente, D.L. Roden, Y. Colino-Sanguino, Z. Kikhtyak, N. Farbehi, J.R.W. Conway, N. Sikta, S.R. Oakes, T.R. Cox, S.I. O’Donoghue, P. Timpson, C.J. Ormandy, and D. Gallego-Ortega. 2021. Single-cell transcriptomics reveals involution mimicry during the specification of the basal breast cancer subtype. Cell Rep 35:108945.

Wanaka, A., S.L. Carroll, and J. Milbrandt. 1993. Developmentally regulated expression of pleiotrophin, a novel heparin binding growth factor, in the nervous system of the rat. Brain Res Dev Brain Res 72:133–144.

Wang, H.Q., and J. Wang. 2015. Expression of pleiotrophin in small cell lung cancer. J Biol Regul Homeost Agents 29:175–179.

Wang, X. 2020. Pleiotrophin: Activity and mechanism. Adv Clin Chem 98:51–89.

Warrington, J.A., A. Nair, M. Mahadevappa, and M. Tsyganskaya. 2000. Comparison of human adult and fetal expression and identification of 535 housekeeping/maintenance genes. Physiol Genomics 2:143–147.

Wellstein, A., W.J. Fang, A. Khatri, Y. Lu, S.S. Swain, R.B. Dickson, J. Sasse, A.T. Riegel, and M.E. Lippman. 1992. A heparin-binding growth factor secreted from breast cancer cells homologous to a developmentally regulated cytokine. J Biol Chem 267:2582–2587.

Weng, T., L. Gao, M. Bhaskaran, Y. Guo, D. Gou, J. Narayanaperumal, N.R. Chintagari, K. Zhang, and L. Liu. 2009. Pleiotrophin regulates lung epithelial cell proliferation and differentiation during fetal lung development via beta-catenin and Dlk1. J Biol Chem 284:28021–28032.

Weng, T., and L. Liu. 2010. The role of pleiotrophin and beta-catenin in fetal lung development. Respir Res 11:80.

Wu, M., M. Ma, Z. Tan, H. Zheng, and X. Liu. 2020. Neutrophil: A New Player in Metastatic Cancers. Front Immunol 11:565165.

Wu, S.Z., D.L. Roden, G. Al-Eryani, N. Bartonicek, K. Harvey, A.S. Cazet, C.L. Chan, S. Junankar, M.N. Hui, E.A. Millar, J. Beretov, L. Horvath, A.M. Joshua, P. Stricker, J.S. Wilmott, C. Quek, G.V. Long, R.A. Scolyer, B.Z. Yeung, D. Segara, C. Mak, S. Warrier, J.E. Powell, S. O’Toole, E. Lim, and A. Swarbrick. 2021. Cryopreservation of human cancers conserves tumour heterogeneity for single-cell multi-omics analysis. Genome Med 13:81.

Yanagisawa, H., Y. Komuta, H. Kawano, M. Toyoda, and K. Sango. 2010. Pleiotrophin induces neurite outgrowth and up-regulates growth-associated protein (GAP)-43 mRNA through the ALK/GSK3beta/beta-catenin signaling in developing mouse neurons. Neurosci Res 66:111–116.

Yang, L., Q. Liu, X. Zhang, X. Liu, B. Zhou, J. Chen, D. Huang, J. Li, H. Li, F. Chen, J. Liu, Y. Xing, X. Chen, S. Su, and E. Song. 2020. DNA of neutrophil extracellular traps promotes cancer metastasis via CCDC25. Nature 583:133–138.

Yao, J., L.L. Zhang, X.M. Huang, W.Y. Li, and S.G. Gao. 2017. Pleiotrophin and N-syndecan promote perineural invasion and tumor progression in an orthotopic mouse model of pancreatic cancer. World J Gastroenterol 23:3907–3914.

Yeh, H.J., Y.Y. He, J. Xu, C.Y. Hsu, and T.F. Deuel. 1998. Upregulation of pleiotrophin gene expression in developing microvasculature, macrophages, and astrocytes after acute ischemic brain injury. J Neurosci 18:3699–3707.

